# Three novel genomes broaden the wild side of the *Capsicum* pangenome

**DOI:** 10.1101/2025.06.03.657591

**Authors:** Christina Papastolopoulou, Ronald Nieuwenhuis, Sven Warris, Linda V. Bakker, Jan van Haarst, Jan Cordewener, Thamara Hesselink, Hetty van den Broeck, Willem van Dooijeweert, Hans de Jong, Julapark Chunwongse, Sara Diaz Trivino, Elio Schijlen, Dick de Ridder, Sandra Smit, Sander A. Peters

**Affiliations:** Bioinformatics Group, Wageningen University & Research, Droevendaalsesteeg 1, 6708 PB, Wageningen, The Netherlands; Wageningen Plant Research, cluster Applied Bioinformatics, Wageningen University & Research, Droevendaalsesteeg 1, 6708 PB Wageningen, The Netherlands; Centre for Genetic Resources, Wageningen University & Research, Wageningen, Droevendaalsesteeg 1, 6708 PB Wageningen, The Netherlands; Laboratory of Genetics, Wageningen University & Research, Droevendaalsesteeg 1, 6708 PB, Wageningen, The Netherlands; Center for Agricultural Biotechnology, Kasetsart University, Kamphaeng Saen, Nakhon Pathom 73140, Thailand

**Keywords:** pepper, wild species, pangenomics, *Capsicum annuum*, *Capsicum chacoense*, *Capsicum galapagoense*

## Abstract

This study presents three genome assemblies within the *Capsicum* genus, enabling comprehensive comparative analyses for the *Annuum* and *Baccatum* complexes within the genus. We produced highly continuous assemblies of the nuclear genomes and complete chloroplast assemblies. Subsequent genome annotation identified 34,580 genes in non-pungent *C. annuum* cv. ECW, and 32,704 and 33,994 genes in pungent *C. chacoense* and *C. galapagoense*, respectively. These assemblies, including the first complete genomes for *C. chacoense* and *C. galapagoense*, provide additional genomic resolution within the *Capsicum* genus. The novel genomes were analyzed within a pangenomic framework, integrating 16 *Capsicum* genomes across the *Annuum, Baccatum*, and *Pubescens* complexes. Homology grouping was used to identify core, accessory and unique genes and showed a wide spectrum of genetic diversity, particularly in homology groups exclusive to *C. chacoense* and *C. galapagoense*. Out of 79,267 homology groups identified, 13% were core groups, present in all accessions, corresponding to approximately 30% of core genes per genome. Comparative analyses revealed distinct species and genus specific genomic characteristics. Additionally, we used the graph pangenome to illustrate locus-level exploration by examining the *Pun1* locus associated with capsaicinoid biosynthesis, identifying multiple *Pun1*-like genes including their genomic position and homology information. The integration of these new resources into a dynamic *Capsicum* pangenome framework provides a versatile platform for extracting genetic information relevant to both fundamental research and breeding applications.

## Introduction

Plant comparative genomics, evolutionary studies, and genomics assisted precision breeding benefit from the rapidly growing availability of sequencing data and the development of high-quality genome assemblies. However, these research fields have also highlighted limitations of comparisons against a single reference genome, as plant genomes can contain up to 90% of repetitive sequences and have complex structural variation, even within the species boundary. Pangenomic analyses, comparing all constituent genomes at once rather than each individually against a reference, have therefore recently become mainstream in plant genomics (Schreiber et al., 2024).

Capsicum is an important food crop, with an annual worldwide production and area harvested amounting close to 40 million tones and over 2 million hectares, respectively (https://www.fao.org/faostat/en/#data/QCL). The *Capsicum* genus belongs to the Solanaceae family, including (among others) tomato, potato, eggplant, tobacco, and petunia. *Capsicum* is well known for its nutritional, medicinal, ornamental, and pharmacological values, and is renowned for its flavoring agent capsaicin, the characteristic ingredient of the spice. Its high vitamin A and C content and richness in iron, potassium and magnesium make capsicum an important nutritional crop. Antioxidant, anti-mutagenic, anti-carcinogenic, and immunosuppressive activities have been attributed to capsaicin as well (Saxena et al., 2016). In the *Capsicum* genus, currently 42 *Capsicum* spp. have been identified, which have been divided over two groups based on their chromosome numbers, 2n=24 and 2n=26 respectively - except for *C. annuum* var. *glabriusculum* that has been profiled as a tetraploid (2n=4x=48) (Back et al., 2024; Jarret et al., 2019; Mongkolporn and Taylor, 2011; Pickersgrill, 1977). *Capsicum* genomes are the second largest of the Solanaceae family, with genome sizes between 3-3.9 Gb, of which approximately 80% is comprised of repetitive elements. Major genomic rearrangements, including extensive chromosomal translocations and genomic rearrangements, contribute to a rich genetic diversity in the *Capsicum* species (Sanatombi, 2024).

Phylogenetic relationships within the *Capsicum* genus were studied by Moscone et al. (2007), who proposed the Annuum (*C. annuum, C. chacoense, C. chinense, C. frutescens*, and *C. galapagoense*), Baccatum (*C. baccatum* and *C. praetermissum*), and Pubescens (*C. pubescens, C. eximium* and *C. cardenasii*, and *C. tovarii*) complexes respectively. Among these species, *Capsicum annuum* and the closely related *C. chinense* and *C. frutescens* are used for food production, with *C. annuum* being the most widely used crop. *C. baccatum* and *C. pubescens* are grown for commercial purposes as well, although to a limited extent. Nonetheless, these closely related species are becoming increasingly important for breeding, as they harbor beneficial traits that can be used for crop improvement.

To benefit from the allelic richness and to advance (introgression) breeding for *Capsicum* crop improvement, assessment of genetic diversity and insight in chromosome topology in the *Capsicum* genus is essential. Disclosing the genetic diversity and chromosome synteny and collinearity is not only critical for precision breeding and selection of compatible breeding parents for inter-specific hybridization breeding but is also required to achieve a deeper understanding of the genetic basis of complex traits. However, while genetic diversity has been studied in terms of SNPs and small insertions and deletions (InDels), structural variations (SVs), are still largely undisclosed. In this respect, *de novo* assembly comparisons are preferred over biased read mapping to capture large SVs (Khan et al., 2020). Other approaches to study genetic diversity, synteny and collinearity involve the use of genetic and physical marker information to position and order alleles. However, genetic markers often have discrepant physical and genetic positions in the genome, complicating precise delineation of genetic loci. Furthermore, most genetic maps for *Capsicum* have been constructed using intra- and interspecific populations which have very few genetic markers in common, and sometimes chromosomes are not congruent with intraspecific maps (Lefebvre et al., 2002; Mongkolporn and Taylor, 2011).

The complex genomic diversity within and among *Capsicum* species poses significant challenges for introgressive hybridization breeding. The observed incongruency between linkage maps, such as that between *C. frutescens* and *C. annuum*, highlights the difficulties in predicting genetic outcomes when translocations occur. This genetic variability can lead to unexpected traits and incompatibilities, making it hard for breeders to develop consistent and desirable varieties. Moreover, while recent pangenome studies have identified valuable genetic variations, they still rely on a single reference genome as an anchor for comparative analyses or are limited to gene-based analysis (Lee et al., 2022; Liu et al., 2023; Ou et al., 2018), which can limit the discovery of novel traits and hinder the full exploration of genomic diversity. A reference-free approach could allow for a more comprehensive understanding of the genetic landscape, enabling breeders to tap into the full potential of the *Capsicum* clade.

To advance our understanding of genomic diversity in the *Capsicum* genus, we present three high-quality *de novo* genome sequences, including two wild *Capsicum* species (*C. chacoense* and *C. galapagoense*), complete with detailed structural and functional annotations. We present a pangenome at the genus level, utilizing PanTools, an alignment-free graph-based approach that effectively eliminates reference bias, ensuring a more accurate representation of genetic variation (Jonkheer et al., 2022; Sheikhizadeh et al., 2018; Sheikhizadeh et al., 2016). The pangenome incorporates structural, functional, and phenotypic annotations, allowing researchers to extract valuable biological insights during analysis. Through this comprehensive framework, we analyze genetic diversity across domesticated and wild *Capsicum* species, as well as show examples of using the pangenome to explore loci of interest across the *Capsicum* genomes. This pangenome facilitates the understanding and mining of *Capsicum* diversification but also lays a foundation for advanced breeding initiatives, with future applications in enhancing food security and improving the livelihoods of communities worldwide.

## Results

### Three *de novo Capsicum* genome assemblies

To enhance our understanding of the genetic diversity within the *Capsicum* genus, we aimed to assemble the genomes of three distinct pepper lines: the non-pungent *C. annuum* cv. Early Cal Wonder (ECW) (CGN16883) and the wild and pungent *C. chacoense* (CGN21477, PI260429) and *C. galapagoense* (CGN22208, AC1501). The goal was to generate genomes that enrich the Annuum and Baccatum complexes within the *Capsicum* genus with both cultivated and wild species. These new genomes complement previously established ones (referenced in Chen et al., 2024; Hulse-Kemp et al., 2018; Kim et al., 2014, Kim et al., 2017; Lee et al., 2022; Liao et al., 2022; Liu et al., 2023; Qin et al., 2014; Shirasawa et al., 2023), providing a more comprehensive resource for comparative genomics and evolutionary studies in *Capsicum*.

For *de novo* genome assembly, we combined multiple (sequencing) technologies. This included large-insert sequencing with PacBio Sequel technology, massively parallel Illumina-based linked-read sequencing with 10X Genomics technology, and Bionano-based genome mapping technology (Table S1). These data were used to assemble and subsequently scaffold the resulting contigs up to chromosome-length sequences. For *C. chacoense* and *C. galapagoense* we scaffolded the genomes in large scaffolds with each chromosome being represented by one to three large scaffolds (Figure S1). For *C. annuum* ECW, we further scaffolded the assembly using a genetic map from Hulse-Kemp et al., 2016, resulting in a near chromosome level assembly. Ultimately, we obtained genomes with total lengths ranging from 3.04 to 3.13 Gb and an N50 final scaffold length from 213 Mb to 256 Mb (Figure 1, Table S2). Completeness of the reconstructed genome sequence was assessed by determining the number of BUSCO (Benchmarking Universal Single-Copy Orthologs) genes. We find at least 97.9% out of a total of 5,950 complete single-copy genes in the Solanales odb10 data set (Manni, Berkeley, Seppey & Zdobnov, 2021; Manni, Berkeley, Seppey, Simão, et al., 2021), indicating a high-quality recovery of the *C. annuum*,

**Figure 1:**
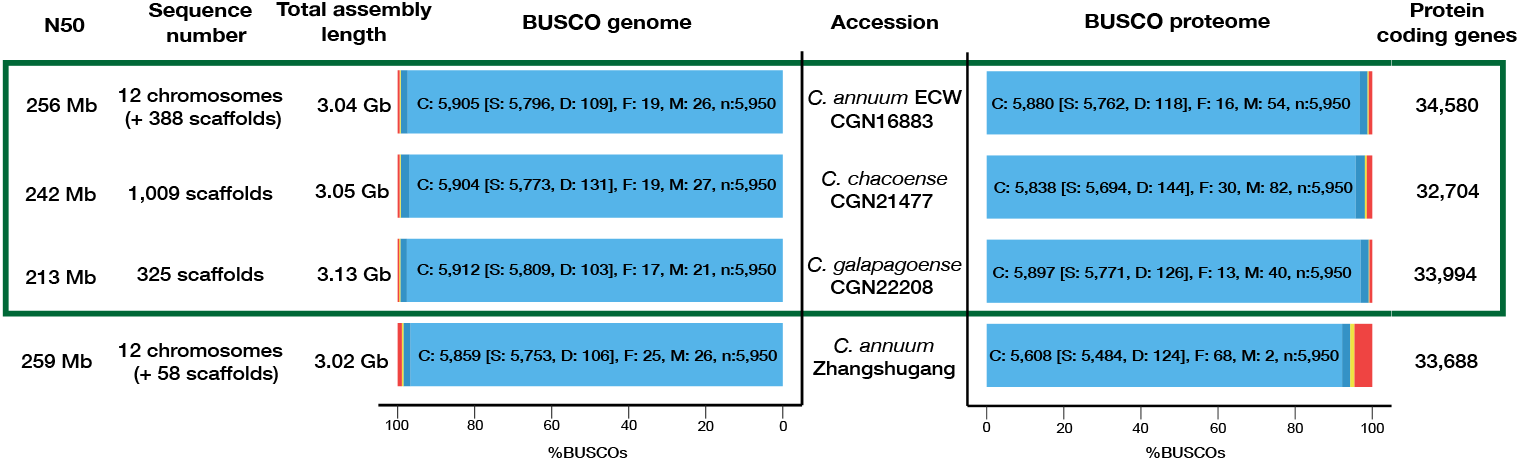
Genome assembly and annotation metrics for the three new *Capsicum* genomes. Genome assembly and structural annotation statistics including BUSCO results for three *de novo Capsicum* genomes, compared to the public *C. annuum* Zhangshugang assembly. The BUSCO score was calculated using the Solanales_odb10 set.

*C. chacoense* and *C. galapagoense* genomes (Figure 1, Table S2). We assessed the completeness of the chromosomes in *C. annuum* ECW with 5,303 markers aligning confidently to the 12 assembled chromosomes, indicating the successful scaffolding and reconstruction of the genome. In chromosomes 1 and 8, the alignment of the markers was inconsistent with the chromosome reconstruction due to a known translocation in the interspecific marker set, while in chromosome 2 a part of the chromosome is not covered by markers (Figure S2). Next to that, we identified nine out of the 24 telomeres in the assembly (Figures S3). All three genomes displayed a repetitive sequence content of 80%, which is similar to previously published *Capsicum* genomes (Chen et al., 2024; Kim et al., 2021, 2017, 2014; Lee et al., 2022; Liao et al., 2022) (Table S3).

In addition to the nuclear genome assemblies, the complete chloroplast genome assemblies (a single contig for the three *Capsicum* species, measuring 156 Kb) demonstrated a high consistency with previously published chloroplast genomes (Figures S4-6). This reinforces the reliability of the genomic data obtained and highlights the stability of chloroplast genome structure across these species.

Next, we combined *ab initio* predicted gene models with evidence-based predictions, annotating 34,580 protein-coding genes for *C. annuum*, 32,704 for *C. chacoense*, and 33,994 for *C. galapagoense*, with 98.3% of protein BUSCOs or more (Figure 1, Table S2). This result indicates that we have achieved a high level of structural annotation completeness.

### Comprehensive *Capsicum* pangenome analysis reveals high genetic diversity and evolutionary relationships across domesticated and wild species

In total, 16 annotated genome assemblies from domesticated and wild *Capsicum* species were selected to construct a *Capsicum*-clade pangenome representing the Annuum, Baccatum, and Pubescens complexes. Specifically, nine *C. annuum* domesticated cultivars were included, along with *C. chinense* PI159236, *C. baccatum* var. *pendulum* PI632928, and the wild *C. annuum* var. *glabriusculum, C. chacoense* CGN21477, *C. galapagoense* CGN22208, *C. baccatum* var. *baccatum* PBC81, and *C. pubescens* Grif 1614, thus covering variation among the three *Capsicum* clades, including wild species, while retaining valuable information from several *C. annuum* accessions (Table S4).

To explore the evolutionary relationship of the newly introduced genomes to the rest of the published *Capsicum* genomes, we estimated genomic distances using two methods. The core-gene approach focuses on the conserved (core) genes shared among the genomes, providing insights into their evolutionary history, while the *k*-mer mash distance approach uses sequence-based comparisons to estimate genomic distances. These methods were chosen to ensure a comprehensive analysis from both gene-based and full genome-based perspectives. The resulting phylogenies were depicted using a bootstrap core SNP tree for the core-gene phylogeny and a clustered heatmap for the *k*-mer mash distance genome relationships (Figures 2, S7). Both methods confirm the existing phylogenies and place *C. annuum* and *C. galapagoense* within the Annuum clade, while *C. chacoense* is placed in the Baccatum clade, which is consistent with previously published results (Carrizo García et al., 2016; D’Agostino et al., 2018; Liu et al., 2023).

**Figure 2:**
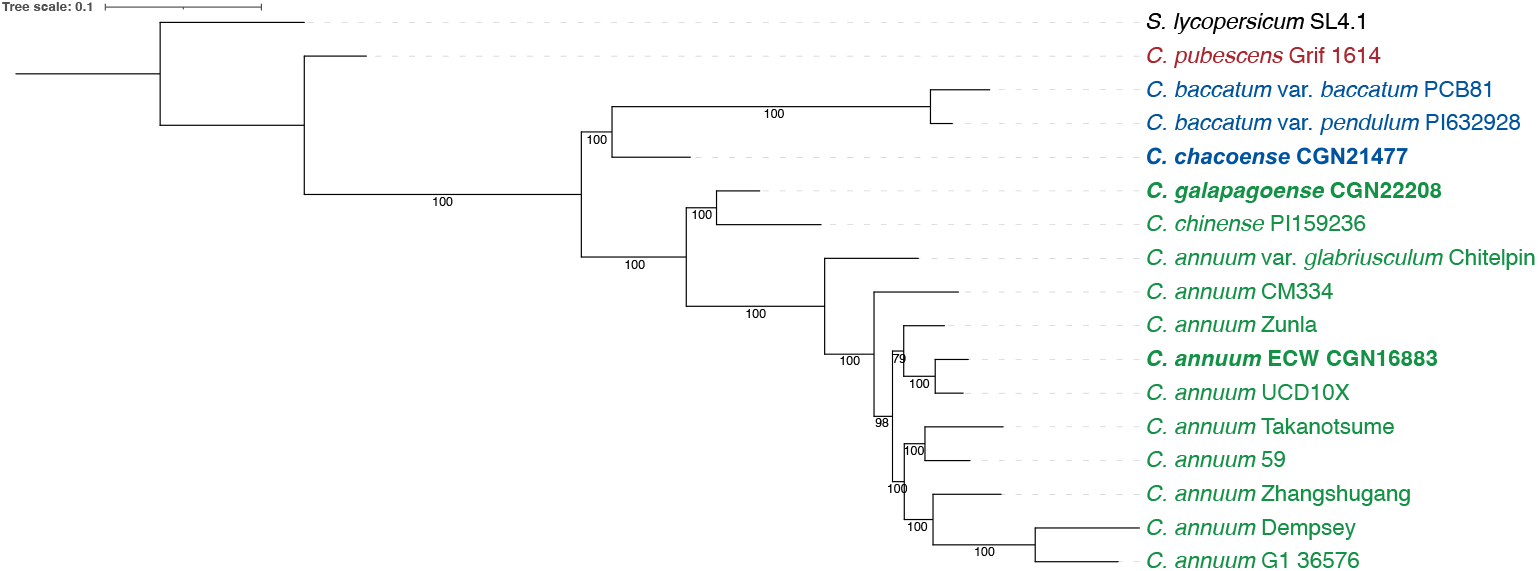
Core phylogeny of *Capsicum* proteomes. A Maximum Likelihood (ML) core SNP tree was constructed based on 6,023 single-copy ortholog groups. Bootstrap values were calculated based on 1,000 iterations, with *S. lycopersicum* SL4.1 serving as the outgroup. The three accessions generated in this study are highlighted in bold, with their respective clades distinguished by color: Annuum (green), Baccatum (blue), and Pubescens (red).

Homology grouping enabled us to study the diversity in genes captured in this pangenome. The 545,775 genes for 16 *Capsicum* accessions were assigned to 79,267 homology groups (HGs), which can be classified based on presence/absence variation in the accession (Figure 3, Table S5). There are 10,368 core groups (13.1%) present in all accessions, corresponding to 29.3-37.6% of core genes per accession (Figure S8). GO enrichment analysis of these core genes revealed their involvement in essential biological processes including transport, photosynthesis, DNA/RNA processes, protein modification/metabolism, and development and defense (Table S6). In contrast to the highly reliable core genes, there are 21,444 unique HGs (only found in one genome), which contain variation in specific processes such as metabolic pathways for specific compounds, developmental processes specific to reproduction and plant defense (Table S7). However, these putative genes (corresponding to 1.4-14.6% per accession) should be treated with care, and the inspection of supporting evidence like function, expression, or experimental validation is highly recommended. More reliable genetic variation is found in the 47,455 (59.9%) accessory groups, occurring in a subset of accessions (56.1-66.3% of genes per accession). GO analysis of these accessory groups showed an enrichment for GO-terms related to defense response, specialized developmental processes, and energy metabolism (Table S8).

**Figure 3:**
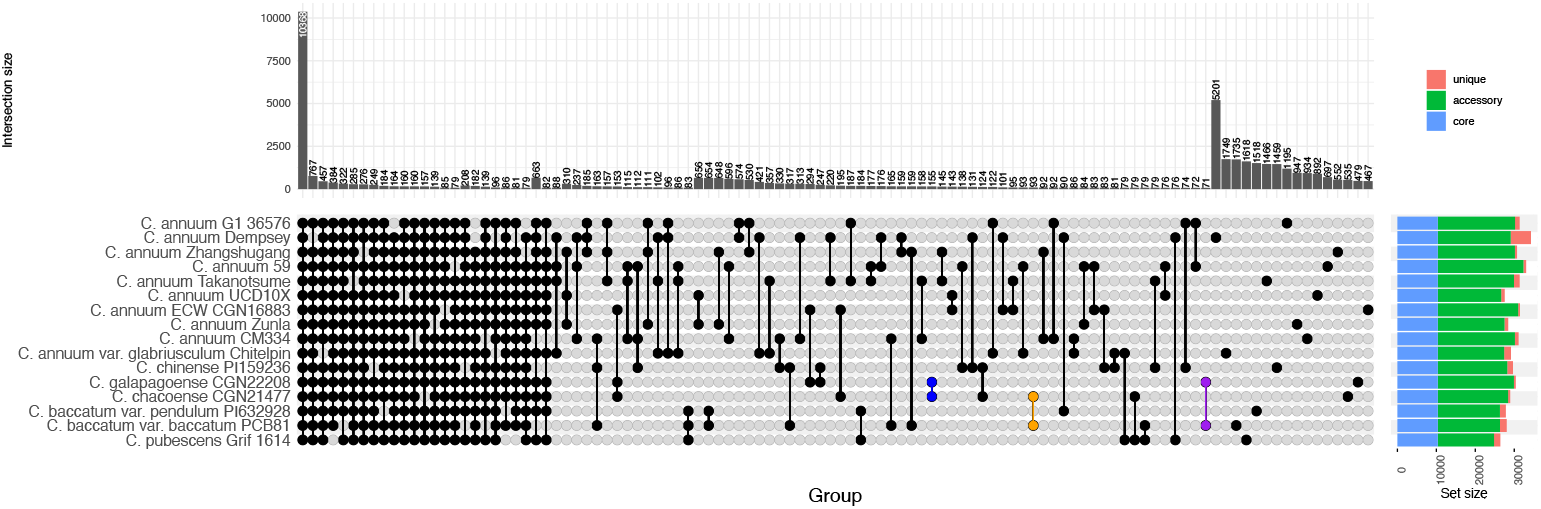
Core, accessory, and unique homology groups in the *Capsicum* pangenome. Upset plot showing the most populated homology groups in the pangenome. The bar plot shows the distribution of the three homology group categories per genome. The unique homology groups between the wild *C. chacoense* and *C. galapagoense, C. chacoense* and *C. baccatum* var. *baccatum* and *C. galapagoense* and *C. baccatum* var. *baccatum* are highlighted in blue, orange and purple respectively.

These genes were further subdivided based on their presence/absence pattern in wild and domesticated accessions (as defined in Table S4). Groups specific to domesticated genomes showed enrichment for terms related to growth and yield, adaptation to cultivation conditions, stress tolerance, plant architecture, and changes in reproductive development (Table S9). In contrast, the 5,008 groups specific to wild species showed enrichment for more fundamental survival-oriented processes with emphasis on defense response and energy management (Table S10). Interestingly, 510 accessory groups of genes are only found in a combination of the four wild accessions, of which 155 apparently are exclusive to both *C. chacoense* and *C. galapagoense*, 93 to *C. chacoense* and *C. baccatum* var. *baccatum*, and 71 to *C. galapagoense* and *C. baccatum* var. *baccatum* genomes (Figure 3). Notably, *C. chacoense* and *C. galapagoense* contain 1,169 unique and accessory groups specific to themselves, with GO terms enriched for binding activities and defense response (Table S11). Considering that domestication in general selects for traits beneficial to humans (e.g. growth, development, reproduction), while wild species maintain a broader range for traits related to survival (e.g. defense and stress tolerance), the observed differences in enrichment therefore possibly reflect selection of human preferred traits historically targeted by *Capsicum* domestication. These results highlight unique genetic features of the novel genomes, strengthening the wild and domesticated parts of the pangenome with potentially interesting genes/alleles for breeding.

We used homology groups as a statistic to assess the saturation of our pangenome. By progressively adding genomes and subsequently analyzing the number of core, accessory, and unique genes we can use the homology groups as a proxy for “new” genes. We plotted the number of core homology groups (present in all genomes so far) as a function of genome number. The declining and stabilizing core curve is clearly plateauing (Figure S8), indicating the level of saturation for the core genes of the *Capsicum* genus pangenome.

## Chromosome structural variations reveal evolutionary divergence in *Capsicum*

Previous comparative sequence alignment studies revealed species- and genus-specific collinearity breaks in the *Solanaceae* clade. These synteny breaks between tomato, pepper, and potato were validated through comparative FISH analyses, ruling out the possibility that collinearity disruptions were due to assembly errors (Peters et al., 2012).

In this study, we analyzed the chromosomes structure of *C. galapagoense, C. chacoense, C. annuum* ECW, *C. pubescens* and *C. baccatum* genomes, using comparative sequence alignment with the *S. lycopersicum* cv. Heinz reference genome (Su et al., 2021). Dot plot comparison (Figures S9-12) revealed similar topologies among these five *Capsicum* genomes, aligning with previously observed patterns in chromosomes 2L, 6S, 10L, and 11L of *C. annuum*. These findings support the existence of genus-specific synteny breaks between *Capsicum* accessions in this study and *S. lycopersicum*, while these structural rearrangements apparently are shared between *Capsicum* and potato (Peters et al., 2012). This indicates that the observed synteny breaks in chromosome 2L are specific to the tomato clade and arose after the split of tomato from the common ancestor of *Capsicum* and potato (Peters et al., 2012). Beyond these clade-specific patterns, we observed additional species-specific structural variations. *C. chacoense* exhibited two minor inversions in chromosome 2L compared to *C. galapagoense, C. annuum* ECW, *C. pubescens* and *C. baccatum* which apparently are specific for *C. chacoense*, while *C. pubescens* exhibited additional structural variations in chromosome 11L. Overall, *C. chacoense* and *C. baccatum* showed more frequent structural variation than *C. galapagoense* (Figures S9, S12), consistent with its more distant phylogenetic relationship to *C. annuum* (Figure 2). These findings underscore both clade-specific synteny breaks and species-specific structural variations in the *Capsicum* genus. Moreover, the development of a comprehensive *Capsicum* pangenome facilitates more detailed analysis and comparison of these structural variations across species.

### Copy Number Variation in the *Capsicum* pangenome for the Pun1 locus

Our comprehensive *Capsicum* data set captures rich, previously unexplored genomic variation across three clades, spanning both wild and domesticated accessions, including both pungent and non-pungent varieties (Table S4). The use of this pangenome allows for efficient comparison and analysis of variation in key genes and loci across different genomes, offering insights into genetic diversity at a broader scale. Pungency is a complex polygenic trait that is regulated by multiple quantitative trait loci involving a pathway of at least nine essential biosynthetic genes. As this trait is of major interest in pepper breeding programs, we selected the *Pun1* locus as a showcase to examine genetic diversity across both pungent and non-pungent accessions (Chen et al., 2024; Stewart et al., 2007, Stewart et al., 2005). Several studies established the *Pun1* locus position on chromosome 2 (Chen *et al*., 2024; Stewart *et al*., 2005). Initial examination of structural variations across *Pun1* locus in our pangenome revealed that all accessions show collinear organization, without significant structural polymorphisms (Figure 4).

**Figure 4:**
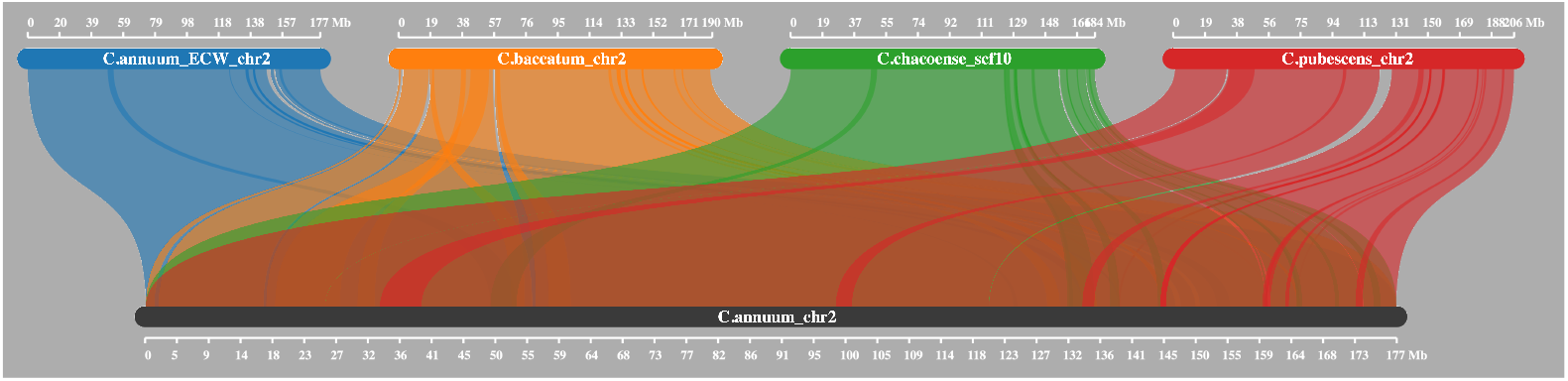
Synteny analysis of chromosome 2 across five *Capsicum* assemblies. Comparison of chromosome 2 syntenic relationships between *C. annuum* cv. Zhangshugang (reference) and four *Capsicum* genomes: *C. annuum* cv. ECW, *C. baccatum* var. *pendulum, C. chacoense*, and *C. pubescens*. The analysis reveals high collinearity in chromosome 2 structure across all examined species.

**Figure 5:**
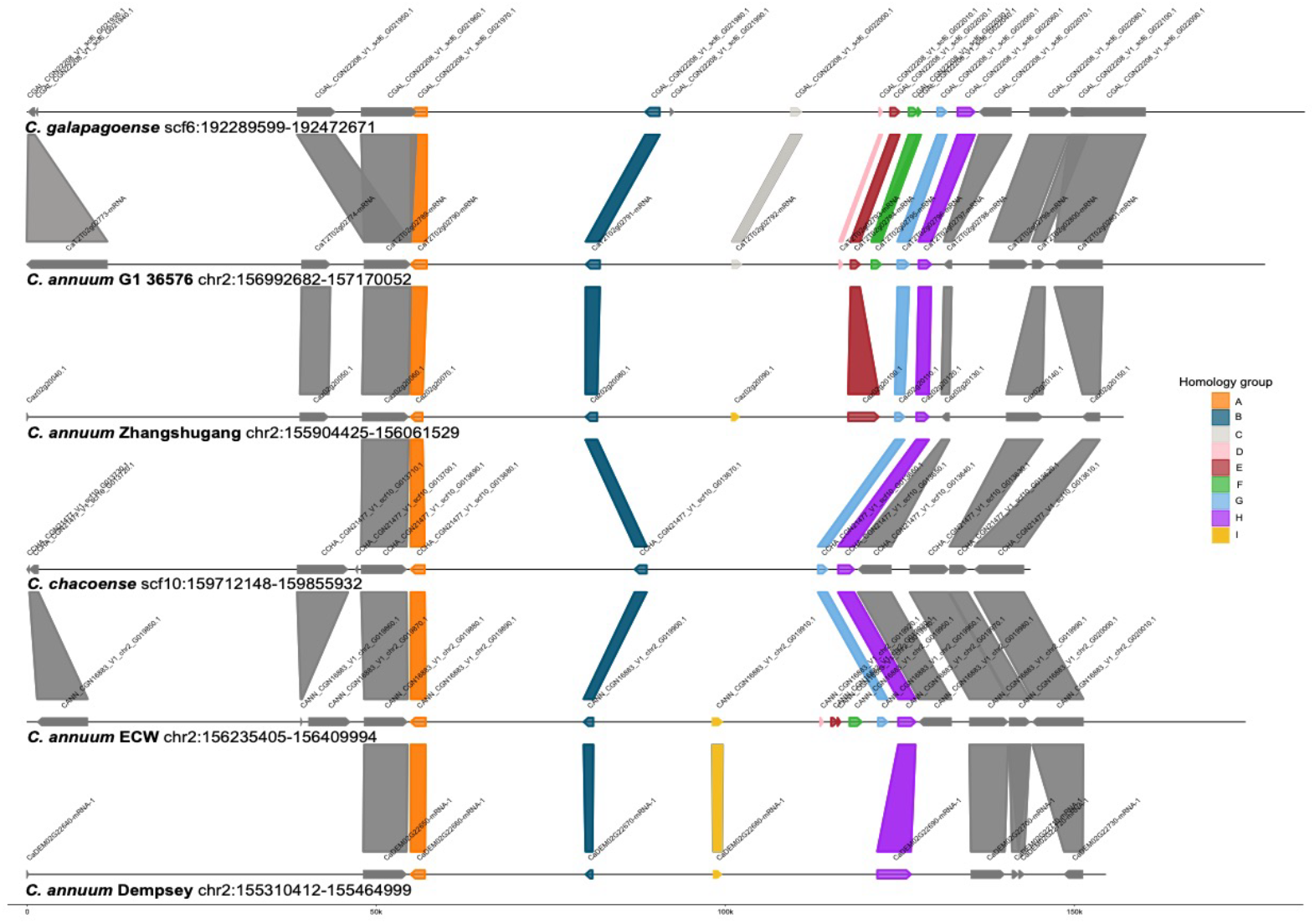
Synteny plot of the *Pun1* region for a subset of the *Capsicum* genomes. mRNAs grouped in the same homology group are connected and mRNAs that had a similarity hit to the *Pun1* gene of *C. annuum* CM334 are marked with the same color per homology group.

Nevertheless, a combination of the homology grouping and a BLASTp search using the *C. annuum* CM334 *Pun1* protein sequence as a query revealed up to nine *Pun1*-like genes, clustering into nine distinct homology groups, with all 16 genomes containing a copy in the same homology group as the query sequence. A maximum likelihood phylogenetic tree confirmed these homology groups and highlighted sequence diversity among the copies. Based on sequence similarity to previous studies, we identified two major homology groups: group A, homologous to capsaicin synthase 1 (CS1); and group B, homologous to capsaicin synthase 2 (CS2). Among our *de novo* genomes, *C. annuum* ECW and *C. galapagoense* contained eight and nine copies respectively, while *C. chacoense* had four copies (Figure S13). However, further analysis is needed to determine which *Pun1*-like copies are functionally active to link these findings to pungency (Chen et al., 2024).

Nonetheless, the above results are illustrative for a graph pangenome-based macro- and micro-synteny analyses, providing critical insights into genomic complexity.

## Discussion

Our study significantly enhances the genomic resources available for the *Capsicum* genus by providing high-quality genome assemblies for three key species: the domesticated *C. annuum* and two wild relatives, *C. chacoense* and *C. galapagoense*. The inclusion of these wild species is significant as they represent two major evolutionary clades within *Capsicum*, harboring traits with high potential for pepper breeding and adaptive studies—traits that were previously inaccessible due to the lack of genomic resources. By leveraging advanced sequencing technologies, we generated a near chromosome-level and two highly contiguous genome assemblies, as reflected by high N50 contig lengths and the BUSCO results. These assemblies and their detailed annotations represent a robust foundation for comparative genomic analyses and advance our understanding of genetic diversity within the *Capsicum* genus opening new avenues for breeding applications and phylogenetic research, providing crucial insights into the evolutionary relationships and trait variability across *Capsicum* species. To further explore genomic diversity, we constructed a *Capsicum*-clade pangenome, incorporating 16 genome assemblies along with their coding genes and their functions spanning both domesticated and wild species. This dataset includes nine *C. annuum* cultivars alongside a diverse array of accessions. By integrating these additional genomes, we captured significant genomic diversity and complexity. Compared to previous studies, such as Lee et al. (2022), which focused primarily on a core *C. annuum* collection with 11 additional accessions, our approach resulted in a substantially larger number of homology groups and a significant reduction in core groups—from 20,000 to just 10,000 (Figure 2). Furthermore, the number of core homology groups for the different genome combinations within in the *Capsicum* pan-proteome clearly levels off, while the dispensable (unique and accessory) still increases (Figure S8), suggesting that the genus-level *Capsicum* pangenome is fully capturing the core genes of the genus but more genomes are needed to capture all the variation. These results provide deeper insights into allelic diversity at the genus level, offering a valuable resource for identifying trait-associated genes and informing breeding strategies. Furthermore, the resources presented here will be supportive to future evolution and genetic diversity studies as well as the development of advanced breeding applications. The use of the pangenome enables us to efficiently analyze and compare variations in key genes in loci of interest across the genomes, offering valuable insights into genetic diversity on a broader scale. For example, pungency, a complex polygenic trait regulated by multiple quantitative trait loci and involving at least nine essential biosynthetic genes, is of key interest in pepper breeding programs. Here, we focused on the *Pun1* locus, a critical determinant of pungency, and examined its genetic diversity across both pungent and non-pungent accessions (Chen et al., 2024; Stewart et al., 2007, Stewart et al., 2005). We used the *Pun1* protein from accession CM334 as a query to identify homologs across a range of accessions and the pangenome to extract the relevant genomic information. This example highlights the interconnection between pangenomes, synteny analysis, and gene model correction.

While we identified highly similar proteins in 16 accessions, we revealed copy number variation and distinct differences between *Pun1* gene copies. Our attention was drawn to the *Pun1* gene in *C. chacoense*, which at first had an unusually large size. Only after inspection of aligned full-length transcriptome data, the *Pun1* gene annotation for *C. chacoense* could be corrected. This example illustrates that pangenomic approaches do not only provide insight into gene arrangements and CNV but also may draw attention to discrepancies in gene annotations and/or genomic sequences that need closer inspection and possibly correction.

However, the pungency trait is more complex by essentially being a quantitative trait that is affected by both genetic and environmental factors. Besides *Pun1*, several capsaicinoid biosynthesis pathway genes have been identified, and multiple QTLs have been associated with localization of capsaicin in placental and pericarp tissue. Nonetheless, given the transcriptome data that was included in recent publications (Back et al., 2024; Chen et al., 2024), identification of transcriptionally active *Pun1* copies may provide more context for future presence-absence-variation (PAV) studies and linking the expressions of gene copies to the pungency trait.

Structural variation (SV) analysis can leverage complementary methods, each with unique strengths and limitations. Sequence-based approaches, such as whole-genome sequencing, provide a comprehensive and linear representation of SV topologies, capturing the detailed arrangement and structure of genomic segments. In contrast, BAC-FISH (bacterial artificial chromosome-fluorescence *in situ* hybridization) analysis previously offered direct physical evidence of SVs in *Solanaceae* spp. including *Capsicum* (Peters et al., 2012) by visualizing the spatial arrangement of specific markers. While BAC-FISH effectively confirmed the presence and location of SVs with high confidence, it lacked the resolution to elucidate the complete topology of the SV segments located between markers. Conversely, the sequence comparison in this study provides additional indirect evidence of genus and species-specific SVs, though it excels at reconstructing their detailed structural organization. Together, these methods provide complementary insights, with BAC-FISH validating SV presence and sequence analysis elucidating its topology in finer detail. In general, the detection of structural variation (SV) is highly valuable, as it provides both opportunities for enhancing genetic diversity and challenges in managing genomic integrity during introgressive hybridization breeding. Previous data suggest that long-distance intra-chromosomal rearrangements and local gene rearrangements have frequently evolved during speciation in the *Solanaceae* family, with small changes being more prevalent than large-scale differences (Peters et al., 2012). This disruption of gene adjacency and synteny, often due to gene and LTR retrotransposon insertions (Peters et al., 2012), has significant implications for hybridization efforts. A study of 12 *C. annuum* accessions, including sweet, hot, and blocky-type peppers, revealed extensive structural variations (SVs) associated with important traits such as virus resistance, pungency, and fruit morphology (Lee et al., 2022). Notably, transposable elements (TEs) in peppers are highly correlated with SVs and have played a significant role in shaping *Capsicum* genomic diversity (Lee et al., 2022; Peters et al., 2012).

While SVs can enhance genetic diversity and the adaptive potential of hybrids, they can also introduce crossing barriers and linkage drag, which limits the use of germplasm for introgressive hybridization. Understanding the effects of SVs is thus imperative as they can influence breeding efforts. For instance, the suppression of recombination in interspecific hybrids between L-type tomato and S-type *S. lycopersicoides* or *S. sitiens* chromosome 10 is explained by a large paracentric inversion (Chetelat et al., 1997; Pertuzé et al., 2002). In the context of *Capsicum*, the observed large structural variations between our newly assembled genomes and the cultivated *C. annuum* genomes (Figure S1, S9) may explain the compatible and incompatible crossing outcomes between *C. annuum, C. frutescens*, and *C. chinense* (Hazarika et al., 2023). Moreover, we achieved a significant advancement enabled by the newly provided genomic resources, in particular the ability to identify species-and genus-specific structural variations (SVs), which can uncover unique genetic adaptations and further inform targeted breeding strategies (Figure S9-S12). Such insights pave the way for leveraging species-specific traits in breeding programs, enhancing the precision and effectiveness of introgressive hybridization.

In summary, insights from our pangenome analysis, including the three new *Capsicum* genomes, provide valuable resources for elucidating genome content and organization. This understanding aids in evaluating collinearity, synteny, and linkage at both the chromosome and gene levels.

## Materials and Methods

### Plant material

Seeds of three different *Capsicum* lines (*C. annuum* Early Cal Wonder (CGN16883), *C. chacoense* PI260429, *C. galapagoense* AC150) were obtained from Centre for Genetic Resources, the Netherlands (CGN) and were sown, germinated, and grown under standard greenhouse conditions at Wageningen University and Research.

### DNA extraction and PacBio sequencing

Plant leaves were harvested, snap frozen in liquid nitrogen and stored at -80°C. DNA isolation was done using 2 to 3 gram per extraction following standard CTAB based protocols. DNA quantity and purity was analyzed by Qubit (Invitrogen), OD values (NanoDrop) and Fragment Analyzer (Agilent). Obtained gDNA was used for two different SMRTbell library protocols (normal and express kit) without initial DNA shearing. Of each line, fifteen microgram of DNA was used for library preparation using SMRTbell template prep kit 1.0 according to manufacturer’s protocol (PacBio; Procedure and Checklist Preparing > 30-kb SMRTbell libraries using Megaruptor Shearing and BluePippin Size selection on PacBio RSII and Sequel systems). Resulting libraries were used for SMRTbell-polymerase complexing using Sequel Binding kit 2.0/2.1, sequencing primer v3/v4 and sequencing on multiple SMRT cells on a PacBio Sequel instrument. All sequencing of these ‘normal’ SMRTbell libraries was done with approximately 5 pM on plate loading, using sequencing kit v2.1, 120 minutes immobilization time and 600 minutes movie time per SMRT cell. In parallel, SMRTbell libraries were made using the SMRTbell Express Template Prep Kit (PacBio) following the Procedure and Checklist Preparing Greater than 15 Kb Libraries Using SMRTbell Express Template Preparation kit 2. Express SMRTbell libraries were subjected to DNA polymerase complexing using Sequel binding kit v2.1/3.0, Sequel Polymerase 3.0, sequencing primer v4 and sequencing on multiple SMRT cells on a PacBio Sequel instrument. All sequencing of these ‘express’ SMRTbell libraries was done with approximately 10 to 20 pM on plate loading, using sequencing kit 2.1/3.0, 120 minutes immobilization time and 600 minutes movie time per SMRT. We achieved a sequence coverage of 54x, 52x and 75x for the *C. annuum, C. chacoense*, and *C. galapagoense* reference genomes respectively, given an expected size of 3.5 Gb for each genome (Table S1; Moscone et al., 2007).

### *Capsicum* genome sequencing, 10X Genomics Linked-Reads

Frozen leaf material of all three *Capsicum* lines was used for nuclei and ultra-high molecular weight DNA extraction following Bionano Prep Plant Tissue DNA Isolation Base Protocol. Of each line, one nanogram of ultra-high molecular weight DNA was used for preparation of a linked-read library using Chromium genome library, gel bead and multiplexing kit and a Chromium controller, following the Chromium genome reagent kits user. Linked-read libraries were used for 2×150 nt paired-end sequencing on Illumina HiSeq2500 and NovaSeq 6000 instruments.

### *Capsicum* genome optical mapping

Of all three *Capsicum* lines, UHMW DNA was used for Bionano Genomics optical genome mapping. DNA was processed using Bionano Genomics DLS Kit following Bionano-Prep-Direct-Label-andStain-DLS-Protocol rev D. Labelled DNA was loaded on multiple Saphyr v2 chips using separate flowcells for the individual lines and analyzed on a Bionano Genomics Saphyr instrument. Genome mapping data was further processed for molecule quality reporting, *de novo* assembly and hybrid scaffolding using Bionano Access v1.3 and Bionano Solve v3.3. Sequence-specific labelling of approximately 700 ng genomic DNA and subsequent backbone staining and DNA quantification for Bionano mapping was done using a Direct Label Enzyme (DLE-1) according to the manufacturer protocol 30206F Bionano Prep Direct Label and Stain Protocol (https://bionanogenomics.com/wp-content/uploads/2018/04/30206-BionanoPrep-Direct-Label-and-Stain-DLS-Protocol.pdfcontent/uploads/2018/04/30206-Bionano-Prep-Direct-Label-and-StainDLSProtocol.pdf). Chip loading and real-time analysis was carried out on a Bionano Genomics Saphyr® analyzer according to the manufacturer system guide protocol 30143C (https://bionanogenomics.com/wp-content/uploads/2017/10/30143Saphyr-System-User-Guide.pdf). Molecule quality hybrid scaffold report was carried out using the Bionano Solve™ analysis pipeline (https://bionanogenomics.com/support-page/data-analysis-documentation/).

### *Capsicum* PacBio Isoseq

Different plant tissues (roots, flowers, fruits, and leaves) were collected, snap frozen in liquid nitrogen and stored at -80°C. Of each sampled tissue, 50-70 milligram was used for RNA extraction by Direct-zol RNA miniprep (Zymo Research) with minor modifications. RNA quantity and purity was analyzed by Qubit (Invitrogen), OD values (NanoDrop) and Bioanalyzer (Agilent). Of each line, RNA of fruit, flower and root were equally pooled to 2 micrograms, which was used for subsequent Iso-Seq library preparation according to manufacturer’s protocol (PacBio; Procedure Checklist Iso-Seq™ Template Preparation for Sequel™ Systems). RNA of leaves was used separately for Iso-Seq library preparation following the same procedures. Each RNA sample was split in two separate cDNA reactions to minimize the risk of potential length-based PCR bias during the full-length cDNA synthesis. First strand cDNA synthesis employed the Clontech SMARTer PCR cDNA synthesis kit. Next, cDNA reactions were used for large scale PCR amplification with 16 parallel reactions. PCR amplification of the cDNA was performed with 16 and 20 cycles for mixed (roots, flowers, and fruits) and leaf cDNA templates respectively. Per template, parallel PCR reactions were pooled, purified and two different Ampure XP elution fractions were collected and quantified by Qubit and BioAnalyzer. Collected amplified cDNA fractions were pooled equimolar and approximately 5 pmol amplified cDNA was used for SMRTbell template preparation (SMRTbell template prep kit 1.0). SMRTbell-polymerase complexes were made using Sequel binding kit 3.0, Sequel polymerase 3.0 and Sequel sequencing primer kit v4. SMRTbell complexes were loaded at approximately 7 to 8 pM for sequencing on a PacBio Sequel system using multiple SMRT cells with 120 minutes immobilization, 240 minutes pre-extension, and 1,200 minutes movie time per SMRT cell. Resulting subreads were used for circular consensus (CCS) read construction with PacBio SMRT link v12. Subsequently, CCS reads were clustered into high quality isoforms, which were further collapsed to unique isoforms using the PacBio isoseq v 4.0.0 Bioconda package. Protein coding regions were predicted in the transcripts using GeneMarkS-T (Gabriel et al., 2023).

### Genome *de novo* assembly and scaffolding

Three *Capsicum* lines were *de novo* assembled using the Flye assembler (Kolmogorov et al., 2019). For *C. annuum* and *C. chacoense*, Flye version 2.4 was used, while for *C. galapagoense*, the assembly was performed using commit 786ec5c8c271f1712d3d6bb7420210026afddc22, between versions 2.4.1 and 2.4.2, to manage the size of assembled contigs during consensus and polishing steps. The resulting assemblies were then polished using Racon v1.3.3 and Arrow v2.3.3 using PacBio CLR data (github.com/lbcbsci/racon). Scaffolding was then done using Arcs v1.0.5 and LINKS v1.8.6 with 10X Genomics linked-read data (Warren et al., 2015; Yeo et al., 2017).

Extended scaffolding was done using Bionano Genomics Solve v3.3.1 based on optical mapping data from DLE1 labelled DNA. A *de novo* assembly and a hybrid scaffold were constructed with Bionano Genomics Solve v.3.2.1, using molecules larger than 120kb. After detecting conflict regions with the Bionano Genomics Access Suite (v.1.3.0), we manually inspected the conflict regions using 10X linked-reads mapped to the super-scaffolds. Mapping and visualization of the 10X derived scaffolds was done with Longranger WGA v.2.2.2 (Marks et al., 2019) and Loupe v.2.1.1 respectively. Gap filling was performed using pbjelly v15.8.24 on both the scaffolded and non-scaffolded parts (English et al., 2012). A final round of polishing utilizing Pilon v1.23 and 10X genomics linked Illumina PE reads was performed after gap filling based on a mapping derived from Longranger align v2.2.2 (https://support.10xgenomics.com/genomeexome/software/downloads/latest).

Scaffolding using Allmaps v0.8.12 was done on *C. annuum* for chromosome anchoring using markers from Hulse-Kemp et al., 2016. This interspecific map contains a known translocation between chromosome 1 and chromosome 8, which was taken in account for this during scaffolding.

The resulting *de novo* assemblies underwent optimization and filtration to eliminate taxonomic and organellar contamination. Specifically, contigs and scaffolds longer than 10kb, exhibiting a GC percentage between 25 and 45 and demonstrating a Solanaceae taxonomic hit (attained through Blobtools v1.1.1) along with PacBio CLR read coverage ranging from 10x to 300x for the same accession, were retained (Laetsch and Blaxter, 2017). Addressing taxonomic and organellar contamination as well as any residual adapter sequences within the genomic sequence, as identified by FCS-GX v0.2.3 (database build 09/07/2023) and FCS-adaptor, was also part of the process, involving their removal or hard masking (Astashyn et al., 2023). Furthermore, contigs and scaffolds flagged as exhibiting mitochondrial or chloroplast contamination, as determined by MitoHiFi v3.2 (blast hits against publicly available *C. annuum* organelles that were larger than 30% of the length of the respective public organelle), were excluded from the genomic assembly (Uliano-Silva et al., 2023). The completeness and redundancy of the genome assemblies was assessed with BUSCO v5.4.7 (Manni et al., 2021b, Manni et al., 2021a). Telomeric sequences were identified using the plant telomeric repeat sequence TTTAGGG with tidk v0.2.65 (Brown et al., 2025).

### Chloroplast assembly and annotation

To assemble the chloroplast sequence of each of the three *Capsicum* species, PacBio CLR from each line were aligned to the two public chloroplast assemblies available in NCBI. From these mappings, primarily mapped reads were used as input to the tool ptGAUL v1.0.5 with 250x coverage (Zhou et al., 2023). The resulting chloroplast assemblies were annotated using the CPGAVAS2 web server and their orientation was corrected according to the commonly accepted order of long single copy (LSC) region, first inverted repeat (IR), short single copy (SSC) region and the second inverted repeat (Shi et al., 2019).

### Repeat masking and structural and functional annotation

RepeatModeler with the -LTRstruct parameter was used to construct a *de novo* library of repetitive elements specific to the three *Capsicum* nuclear genomes (Flynn et al., 2020). This custom library, combined with the Repbase library (version 26.10.2018) and the Dfam library of transposable elements (TE) sequence alignments and hidden Markov models (HMMs) v3.7, was utilized in RepeatMasker v4.1.5 (tetools v1.87) to identify and soft mask these repetitive elements within the genome assemblies. The genome assemblies with soft-masked repetitive regions were subsequently used for structural annotation.

Braker3 v3.0.3 was used to structurally annotate the *C. annuum, C. chacoense* and *C. galapagoense de novo* assembled genomes (Brůna et al., 2021; Buchfink et al., 2015; Gabriel et al., 2023, 2021; Gotoh, 2008; Hoff et al., 2016; Iwata and Gotoh, 2012; Kim et al., 2019; Kovaka et al., 2019; Pertea and Pertea, 2020; Stanke et al., 2008; Stanke and Waack, 2003). Publicly available RNA-Seq data from various tissues and treatments were selected as input for Braker3 (SRR10011918, SRR10011923, SRR10011924, SRR11874047, SRR13488410, SRR17837298, SRR17837305, SRR771924, SRR771925, SRR771927, SRR771928, SRR771929, SRR771932, SRR771938, SRR771939, SRR771952, SRR8692596). Additionally, protein sequences from the Viridiplantae odb11 and the proteins of *C. annuum* UCD10Xv1.1 (NCBI RefSeq GCF 002878395.1) were used. Gene models were also predicted with Helixer v0.3.2 using the land plant lineage and default parameters. The final gene models were produced by combining and refining the Braker3 and Helixer gene models with EvidenceModeler (EVM) v2.1.0 (Haas et al., 2008). To enhance accuracy, we integrated assembled transcriptomic data from public RNA Seq libraries, along with assembled Iso-Seq data and the protein mappings from Viridiplantae odb11, UCD10Xv1.1 (NCBI RefSeq GCF 002878395.1) and Zunla (GCF 000710875.1). Finally, UTR regions were predicted, and we modelled alternative splicing using PASA v2.5.3 (Haas et al., 2008). Single exon genes without a homolog in another *Capsicum* genome and incomplete gene models were discarded. During the homology grouping analysis, we identified several inconsistencies in the gene models. Based on expression data validation, we replaced PASA genes that had been incorrectly merged with their corresponding EVM models. We assigned functions to the final set of predicted genes with functional annotation of the predicted gene models using InterProScan v5.66-98.0 (Jones et al., 2014).

### *Capsicum* graph pangenome construction

*Capsicum* genome assemblies and annotations for *C. annuum* Early California Wonder (ECW), *C. annuum* UCD-10X-F1, *C. annuum* CM334, *C. annuum* Dempsey, *C. annuum* Zhangshugang, *C. annuum* Zunla, *C. annuum* Takanotsume, *C. annuum* 59 (using the updated annotation from Shi et al., 2024), *C. annuum* G1-36576, *C. annuum* var. *glabriusculum, C. chinense* PI159236, *C. chacoense* CGN21477, *C. galapagoense* CGN22208, *C. baccatum* var. *pendulum* PI632928, *C. baccatum* var. *baccatum* PBC81, and *C. pubescens* Grif 1614 were collected to build a *Capsicum* clade graph pangenome. From the collected genomes, contigs smaller than 5 Kb were removed, and from the annotations, only the longest isoforms, encoding proteins larger than 30 amino acids were selected using the PanTools QC pipeline (https://github.com/PanUtils/pantools-qc-pipeline). The chromosomes and/or larger scaffolds of the input genomes were oriented based on their longest alignment with the respective chromosomes of *C. annuum* Zhangshugang. PanTools v4.3.1 functions build_pangenome, add_annotations, add_functions and add_phenotypes were used to construct a genus-level pangenome from the selected 16 *Capsicum* genome assemblies, including structural and functional annotations with accompanying phenotype data (Jonkheer et al., 2022; Sheikhizadeh et al., 2018; Sheikhizadeh et al., 2016). Homology grouping parameters were optimized with the optimal_grouping function using the solanales_odb10 lineage to monitor false and true positives and negatives, while grouping of the proteins in homology groups was performed using --relaxation 4. Homology groups were classified using the gene_classification function into core, unique and accessory categories, with further subdivision based on wild and domesticated phenotypes (Table S4).

### Phylogeny

Full genome core SNP phylogeny was constructed by classifying genes as core, accessory or unique using PanTools. For the core SNP phylogeny, a multiple sequence alignment of the single-copy core genes in the pangenome was used to generate a maximum likelihood phylogenetic tree, representing the core genome phylogeny using the PanTools core_phylogeny function. Genome sequence distances between genomes were calculated using Mash v2.3 and the triangle function (Ondov et al., 2016).

### GO term enrichment analysis

Gene ontology analysis for molecular function (MF), cellular component (CC) and biological process (BP) was performed with GOATOOLS with pvalue < 0.05 and fdr correction method Benjamini/Hochberg (Klopfenstein et al., 2018)

### Structural variation analysis

Genome assembly alignments were aligned using minimap2 (Li, 2021, 2018) and alignments were visualized using the R package pafr (https://github.com/dwinter/pafr).

### Region of interest analysis

PanTools v4 commit ID a8d2b4e and extract_homology_groups was used to extract the regions of interest based on gene homology. Homology groups that contain potential gene copies of the genes of interest were identified by performing BLASTP search of the query protein, against the pangenome protein database (threshold >=50% identity and >=25% alignment length of input sequence). The region of interest was visualized using the R package gggenomes (https://github.com/thackl/gggenomes).

## Data availability

The raw sequencing data (PacBio CLR, Illumina paired end), genome assemblies and annotations, are publicly accessible and can be retrieved from the National Center for Biotechnology Information (NCBI) under the project ID PRJNA944626.

## Acknowledgements

We would like to thank Zijiang Yang and Dirk-Jan van Workum for their annotation and pangenomics support. This work was funded by the Top consortium for Knowledge and Innovation (TKI) program with project number LWV2004.

## Notes

### Competing Interest Statement

The authors have declared no competing interest.

