## Supplemental Figures for "Three novel genomes broaden the wild side of the *Capsicum* pangenome"

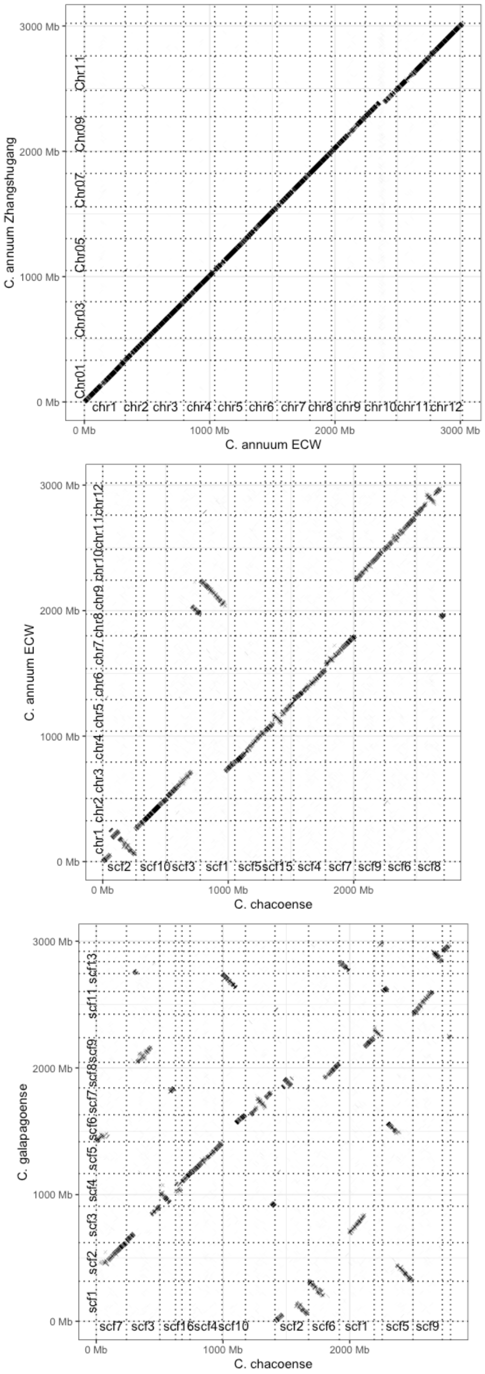


**Figure S1**: Dot plot analyses showing pairwise genomic comparisons between (from top to bottom): A) *C. annuum* cv. Zhangshugang versus *C. annuum* ECW, B) *C. annuum* WUR versus *C. chacoense,* C) *C. galapagoense* versus *C. chacoense.*

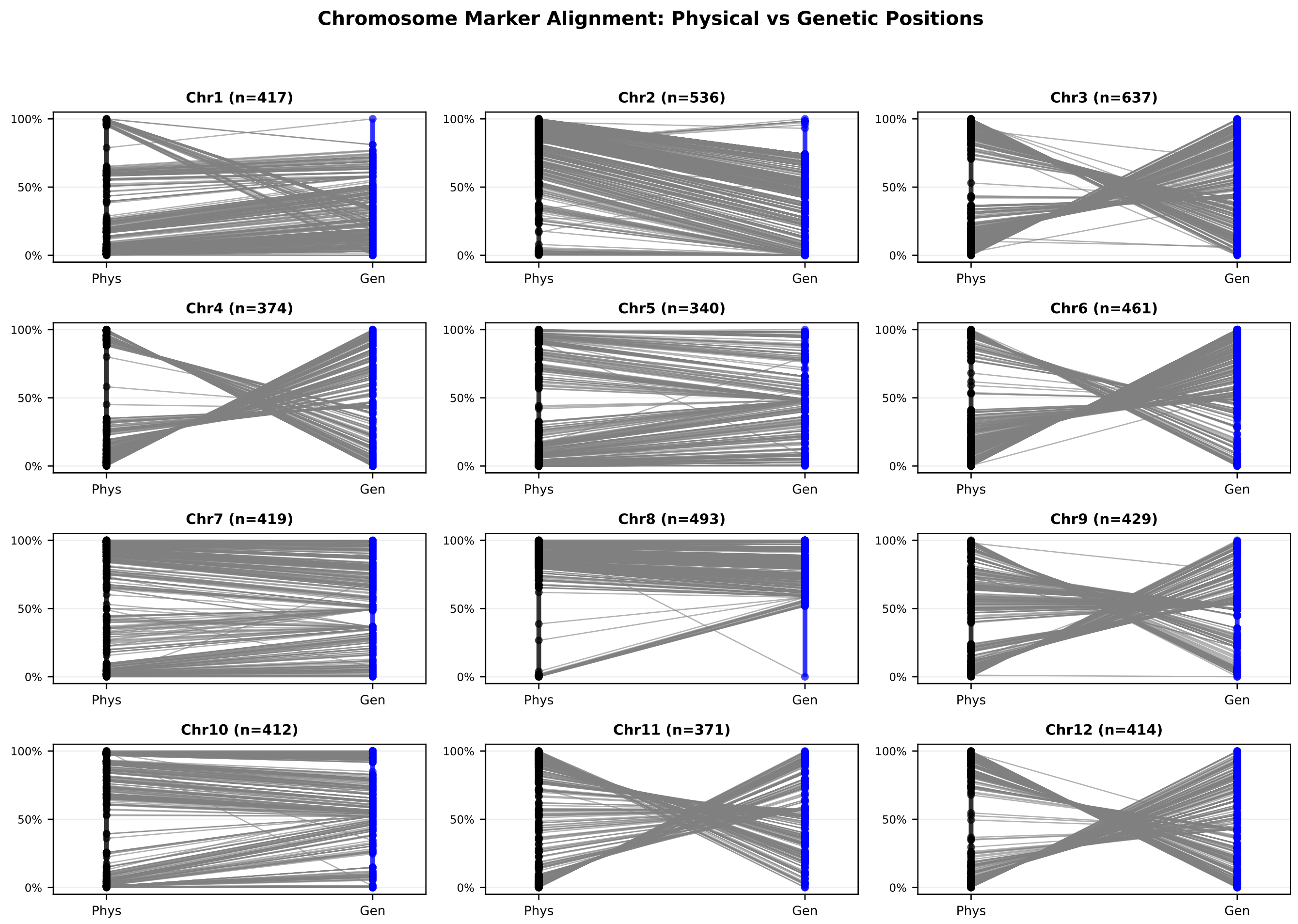


**Figure S2**: Chromosome marker alignment comparison of physical (Phys) and genetic (Gen) positions of the genetic markers from Hulse-Kemp et al., 2016 against the *C. annuum* WUR genome assembly. The chromosome orientation has been decided based on the *C. annuum* Zhangshugang genome assembly.

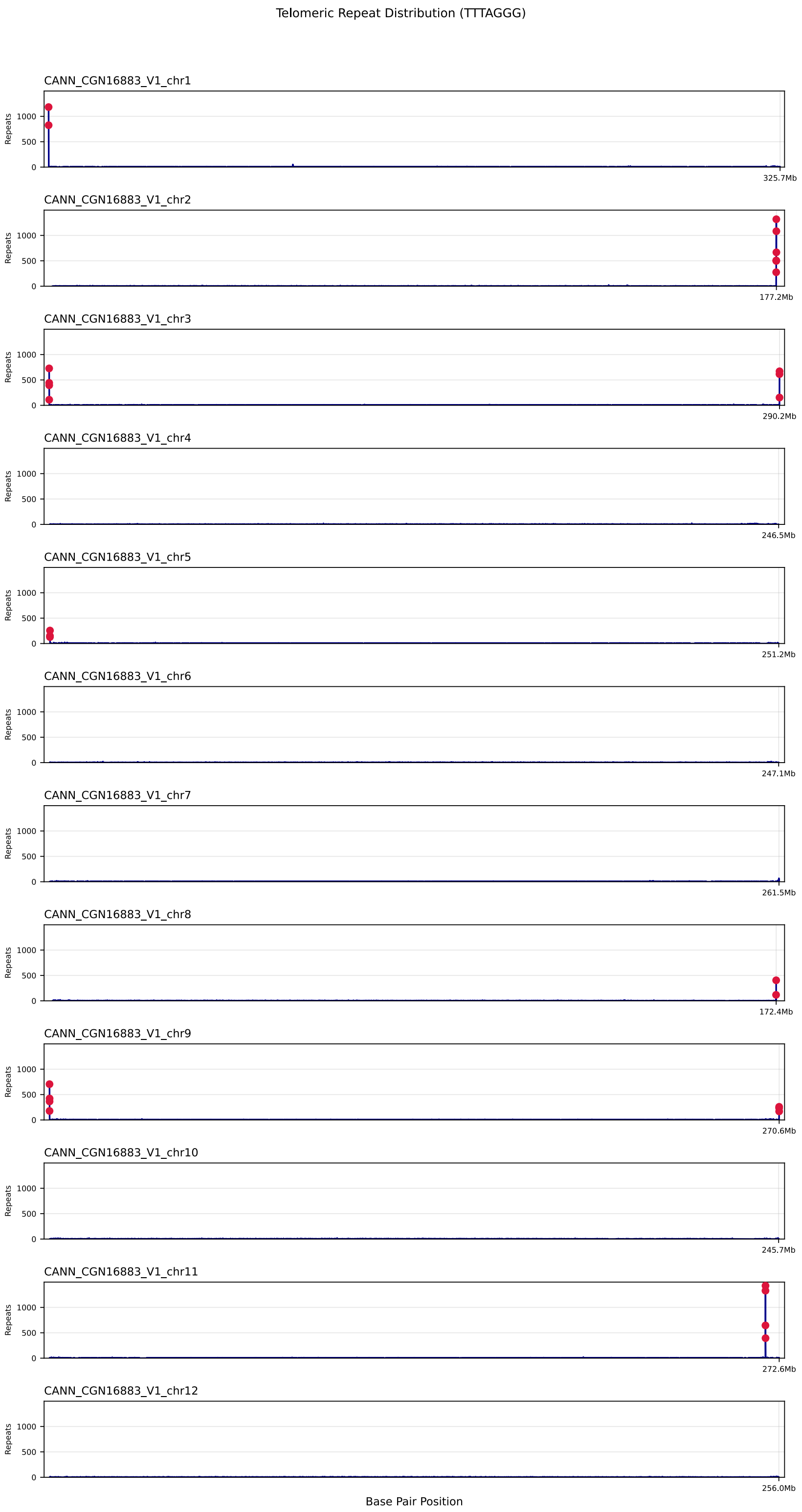


**Figure S3**: Plant telomeric repeat sequence (TTTAGGG) frequency for the 12 chromosomes of the *C. annuum* ECW, showing the presence of nine telomeres. Each red dot marks the presence of at least 100 repeats in a 10kb window.

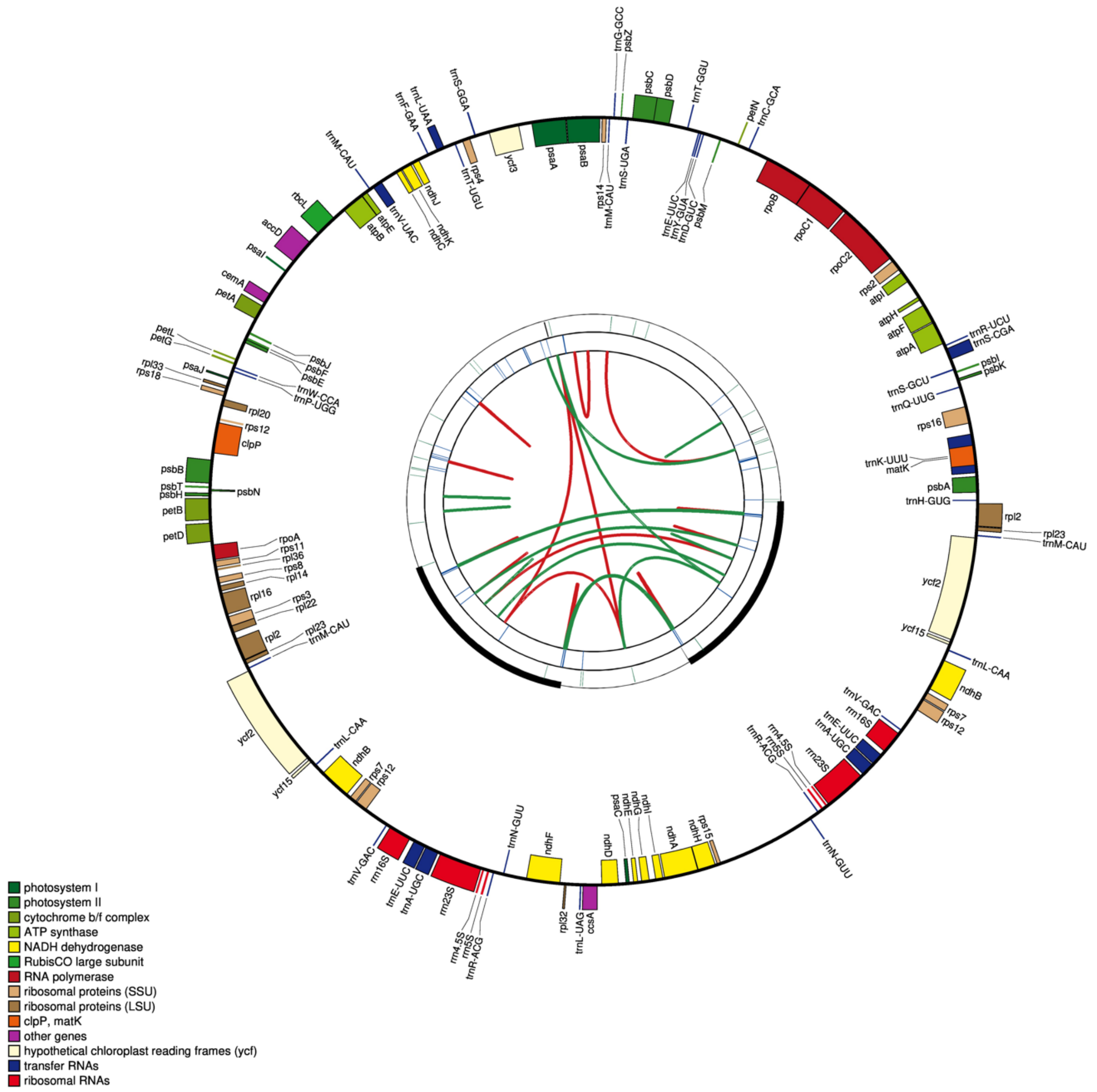


**Figure S4:** *C. annuum* ECW chloroplast annotated by CP2GAVAS.

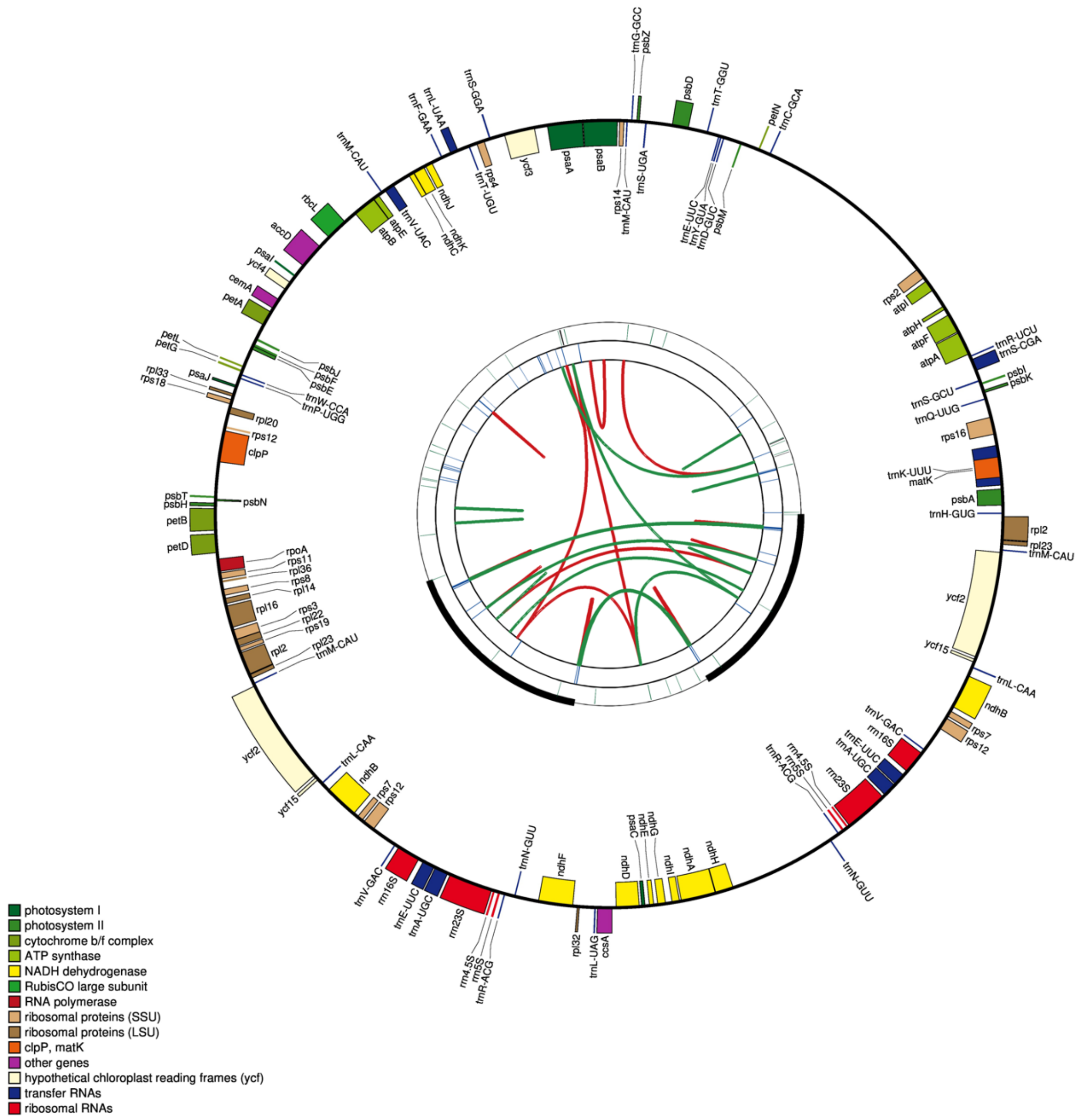


**Figure S5**: *C. chacoense* chloroplast annotated by CP2GAVAS.

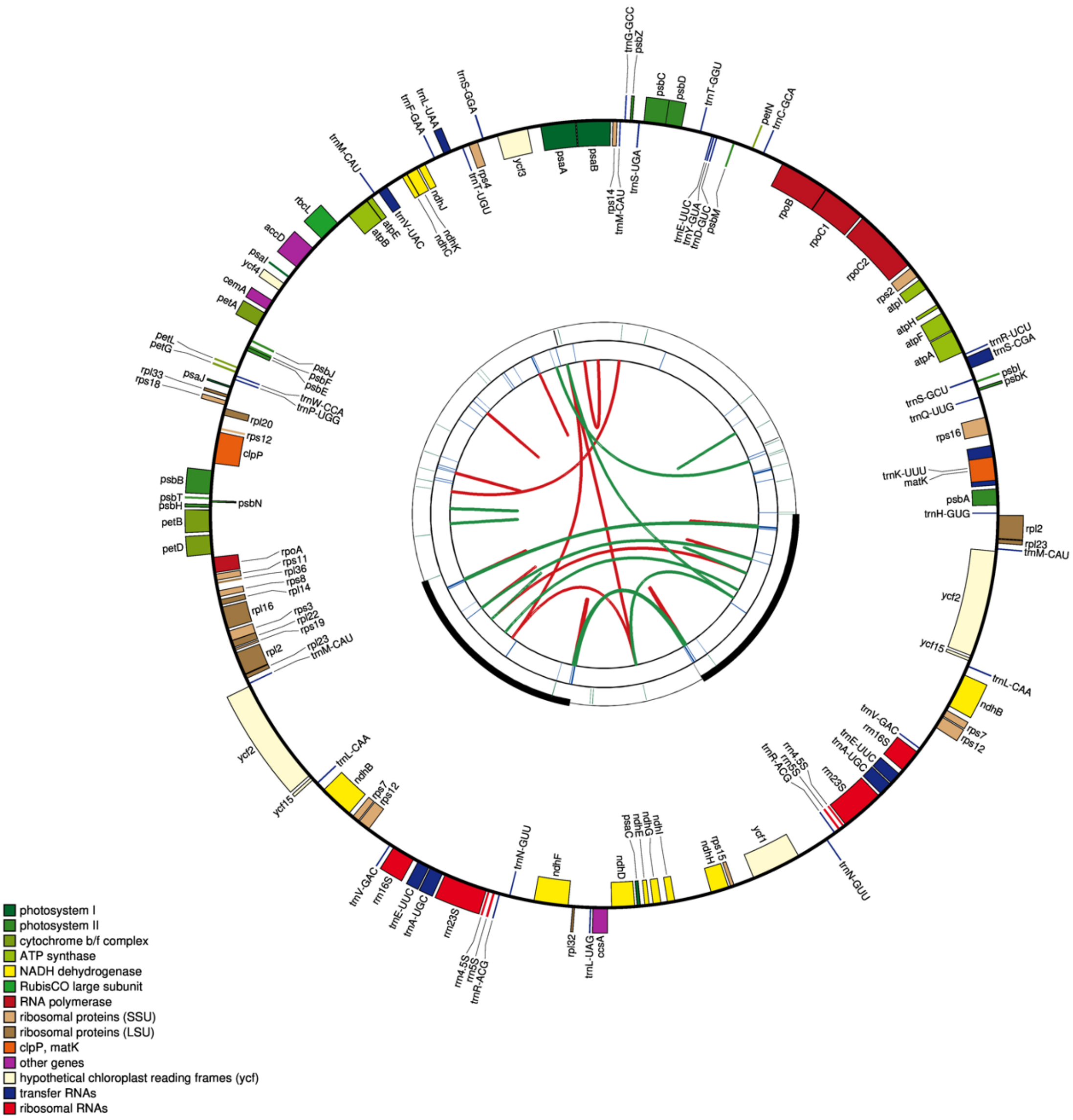


**Figure S6:** *C. galapagoense* chloroplast annotated by CP2GAVAS.

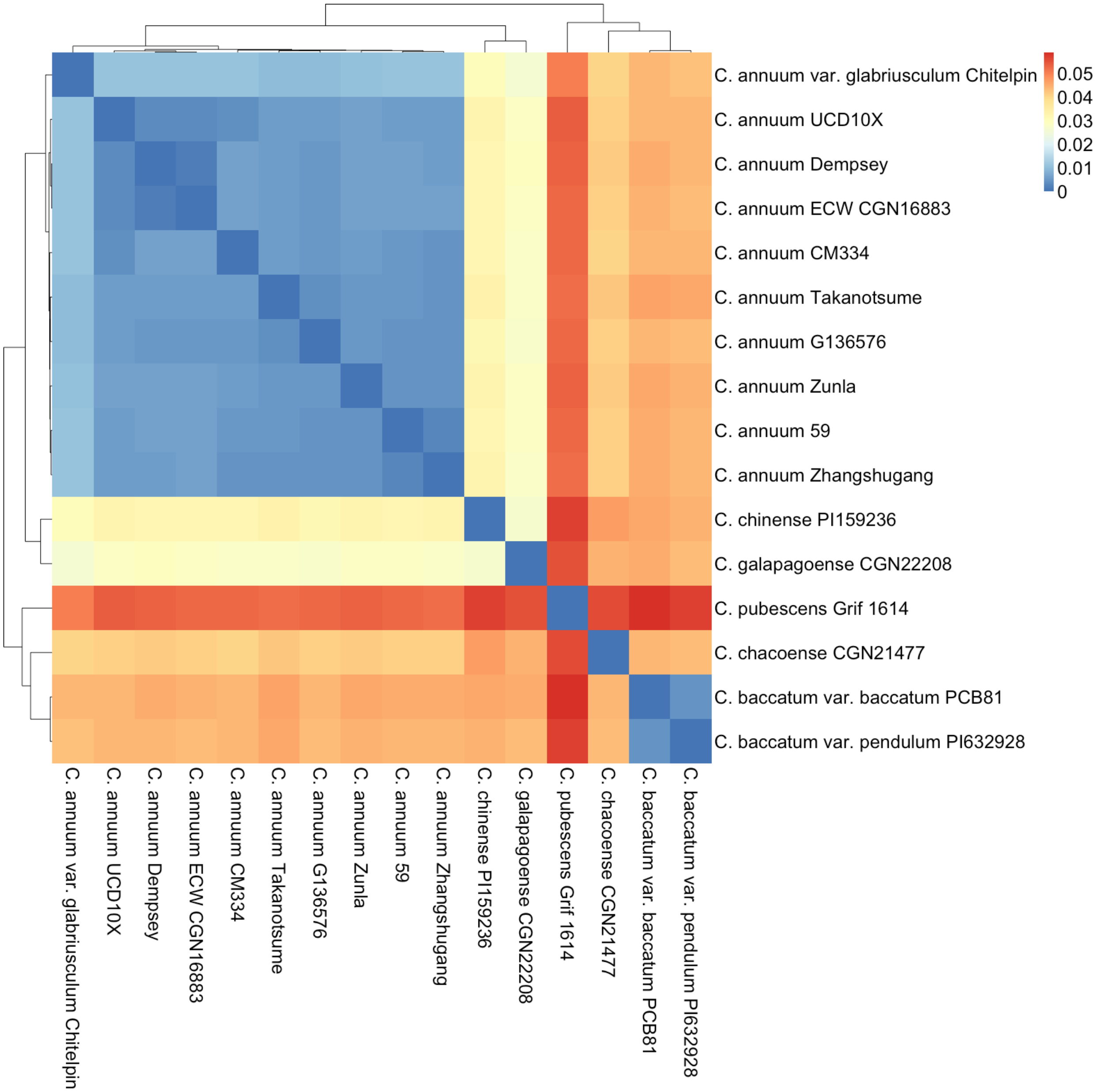


**Figure S7:** Genome similarity for the *Capsicum* genomes. Mash k-mer distance highlights the sequence similarity within the *C. annuum* genomes. The similarity gradually decreases for the other species in the Annuum clade and becomes larger for Pubescens and Baccatum clades.

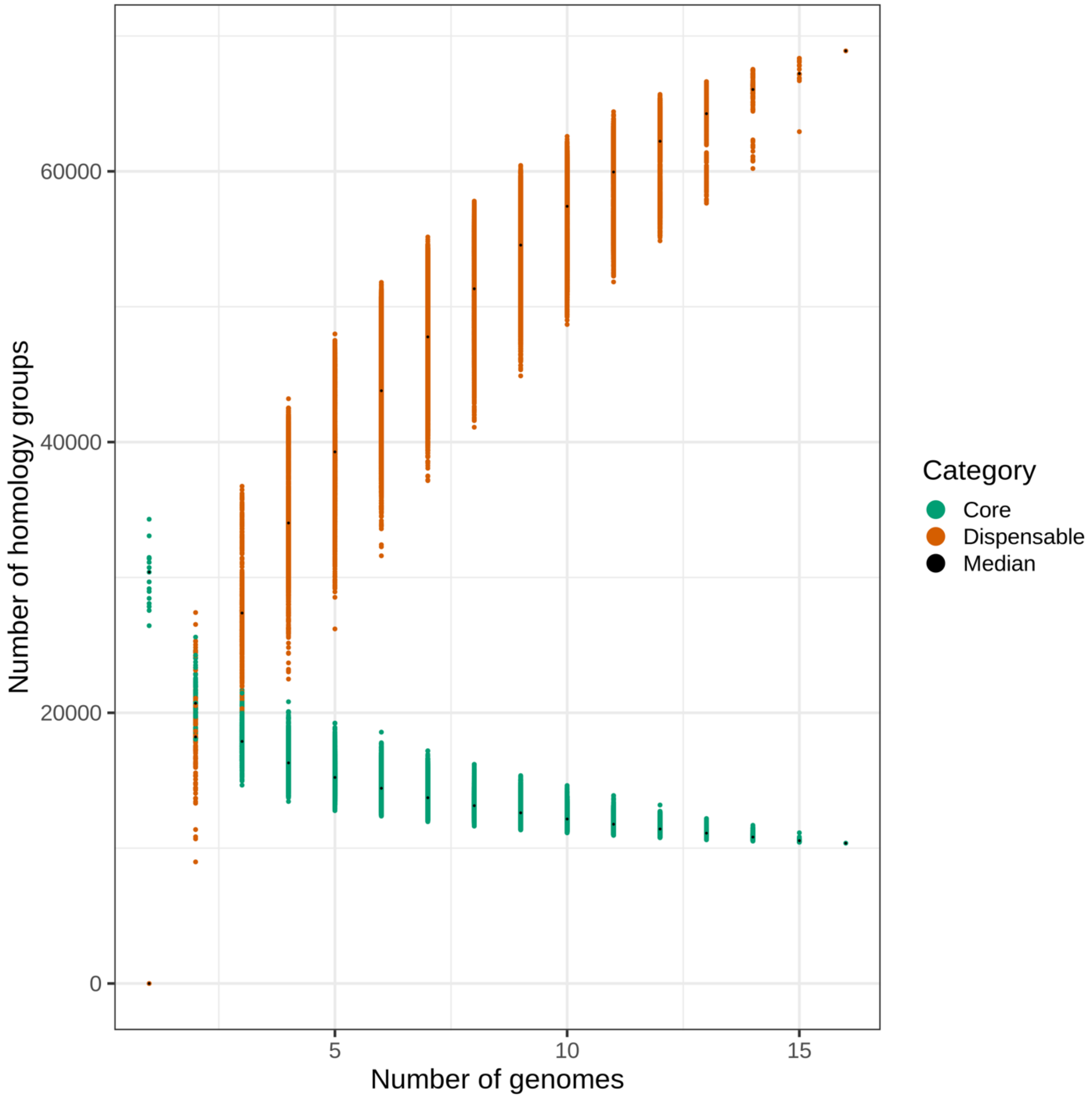


**Figure S8:** *Capsicum* panproteome growth curve. The number of core and dispensable (unique and accessory) homology groups for the different genome combinations within in the *Capsicum* panproteome.

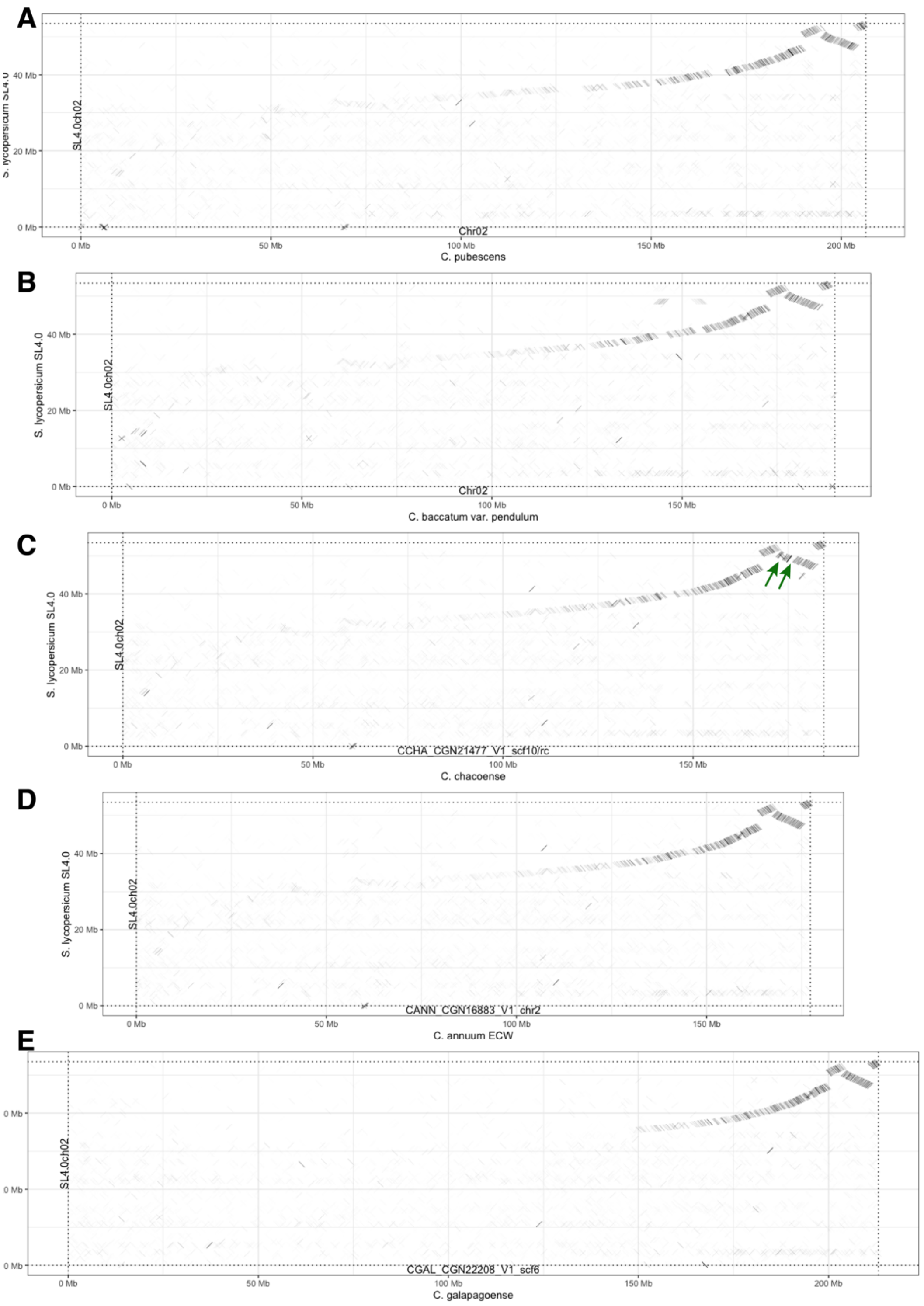


**Figure S9**: Dot plot analyses showing pairwise genomic comparisons of chromosome 2 (from top to bottom): A) *C. pubescens*, B) *C. baccatum* var. *pendulum*, C) *C. chacoense*, D) *C. annuum* ECW, and E) *C. galapagoense* versus *S. lycopersicum* SL4.0.

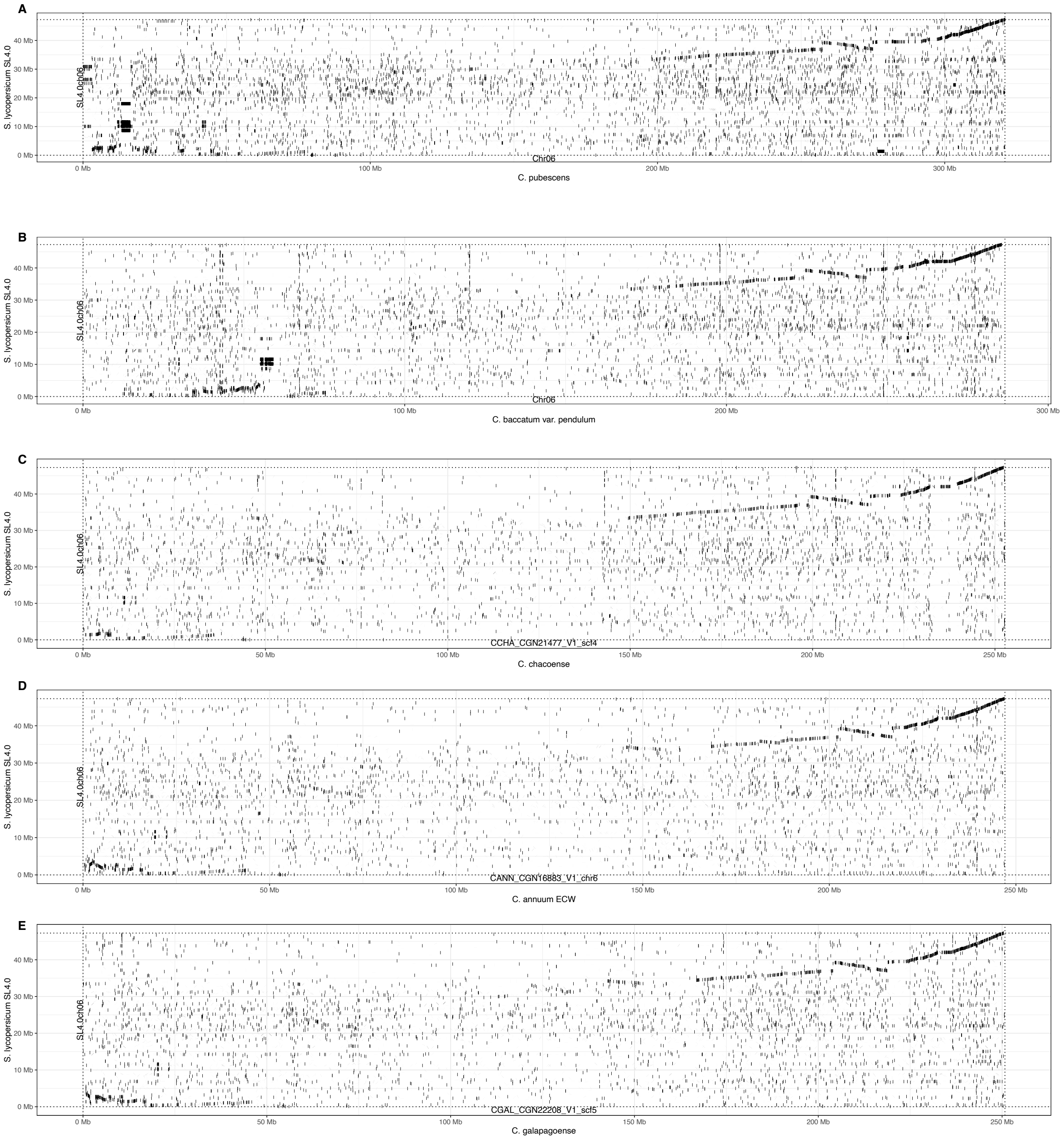


**Figure S10**: Dot plot analyses showing pairwise genomic comparisons of chromosome 6 (from top to bottom): A) *C. pubescens*, B) *C. baccatum* var. *pendulum*, C) *C. chacoense*, D) *C. annuum* ECW, and E) *C. galapagoense* versus *S. lycopersicum* SL4.0.

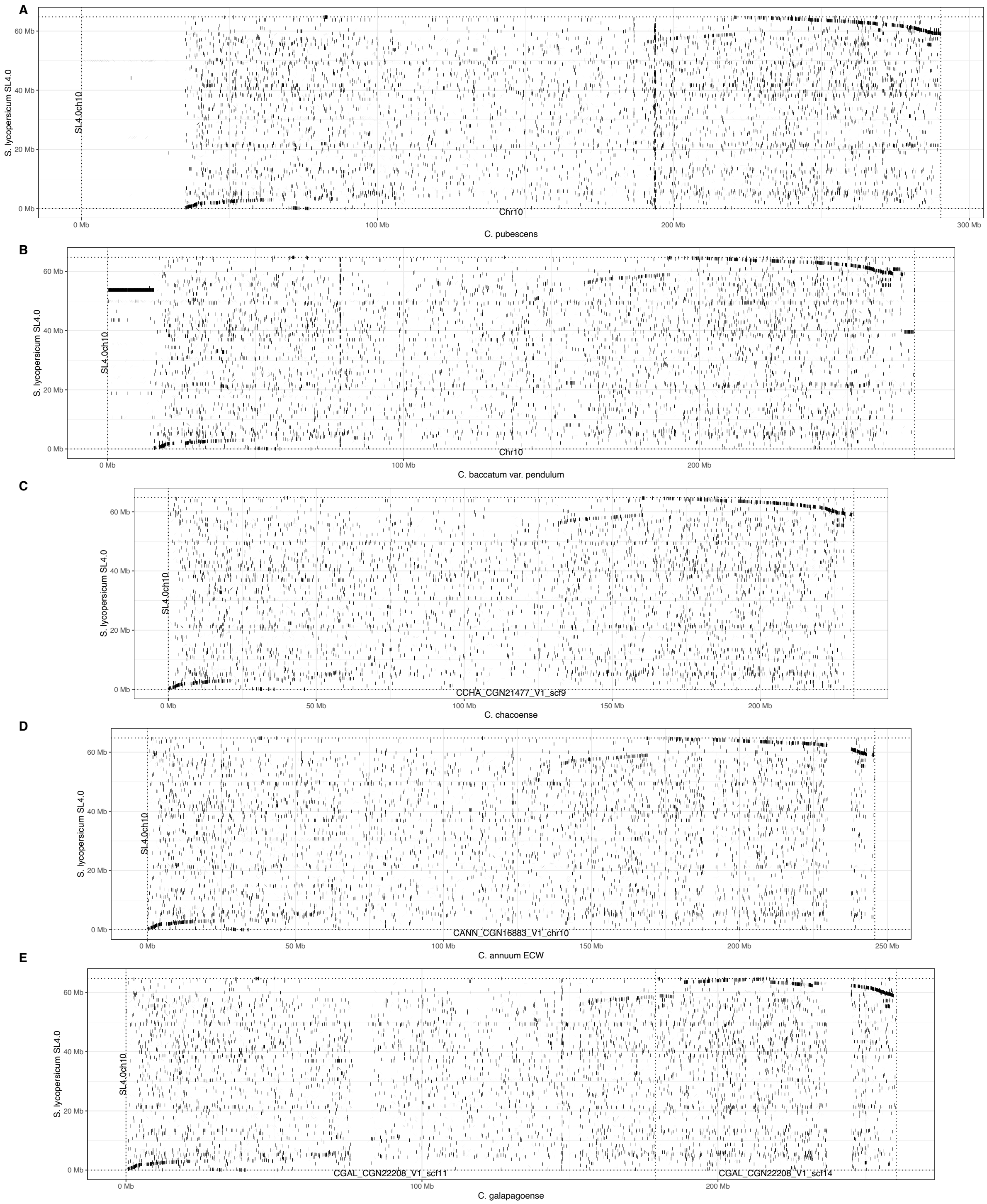


**Figure S11**: Dot plot analyses showing pairwise genomic comparisons of chromosome 10 (from top to bottom): A) *C. pubescens*, B) *C. baccatum* var. *pendulum*, C) *C. chacoense*, D) *C. annuum* ECW, and E) *C. galapagoense* versus *S. lycopersicum* SL4.0.

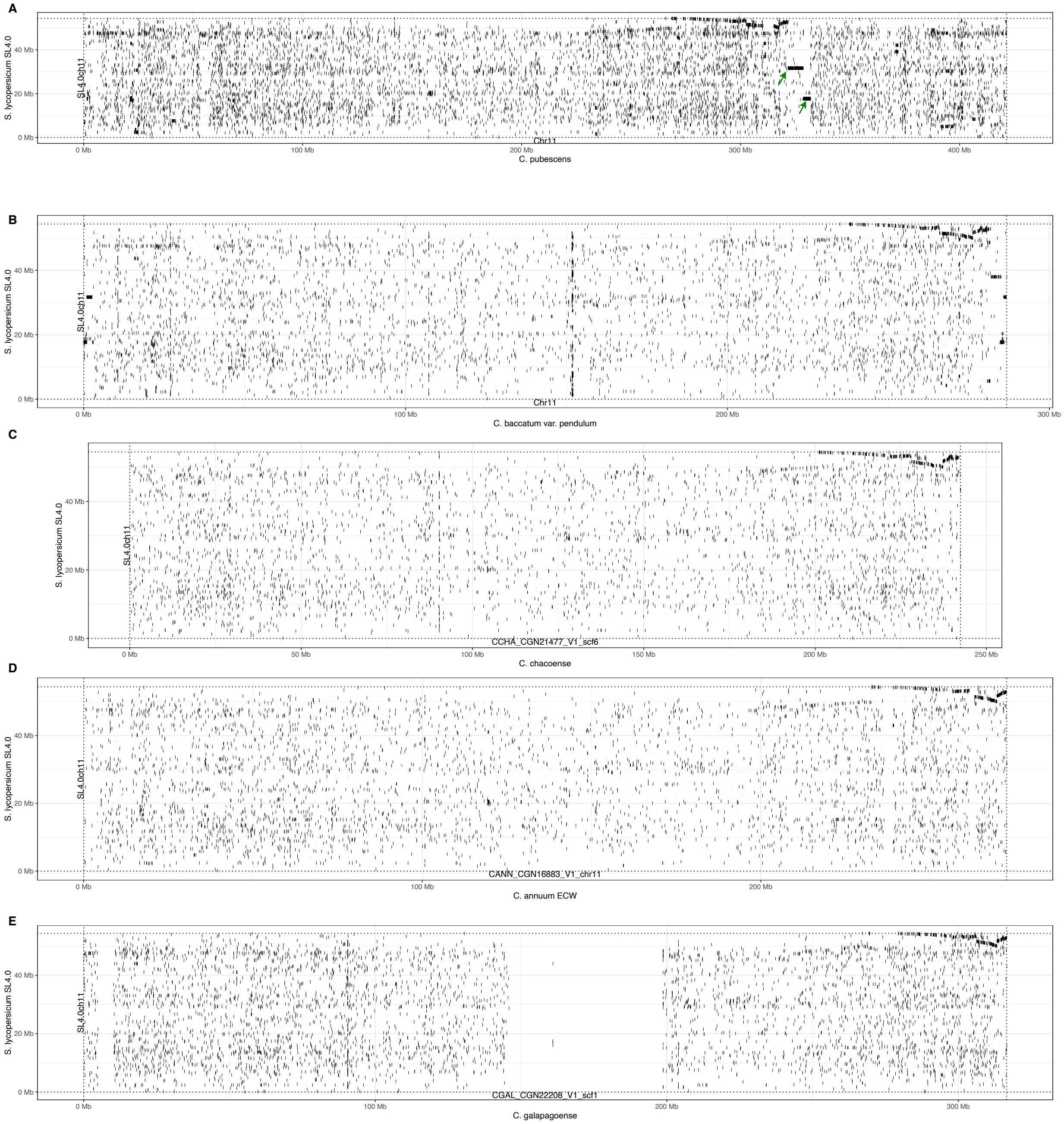


**Figure S12**: Dot plot analyses showing pairwise genomic comparisons of chromosome 11 (from top to bottom): A) *C. pubescens*, B) *C. baccatum* var. *pendulum*, C) *C. chacoense*, D) *C. annuum* ECW, and E) *C. galapagoense* versus *S. lycopersicum* SL4.0.

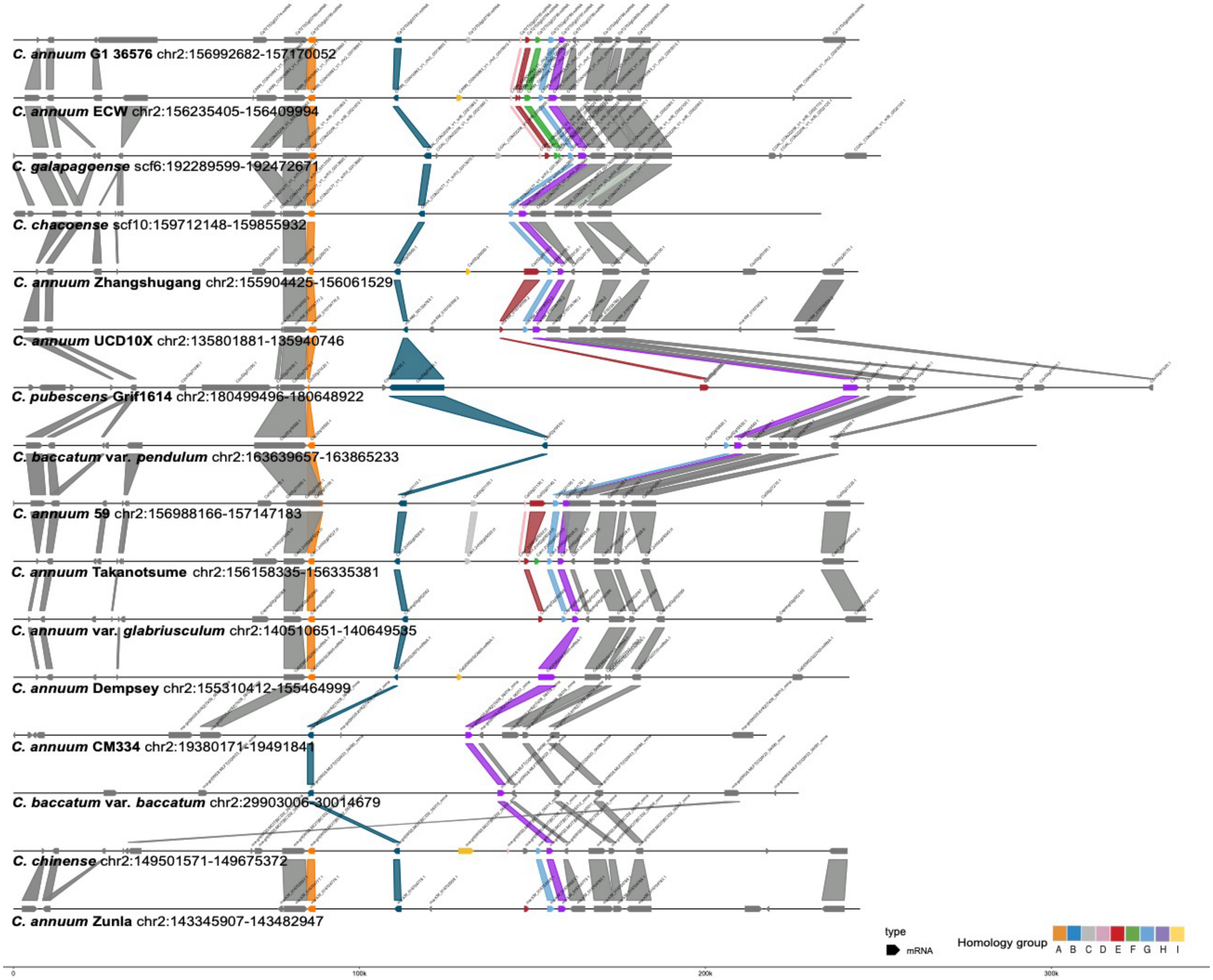


**Figure S13:** *Pun-1* locus for the 16 *Capsicum* genome assemblies. The *Pun-1* gene copies are organized in nine homology groups, ranging from one to nine *Pun1*-like gene copies per genome.

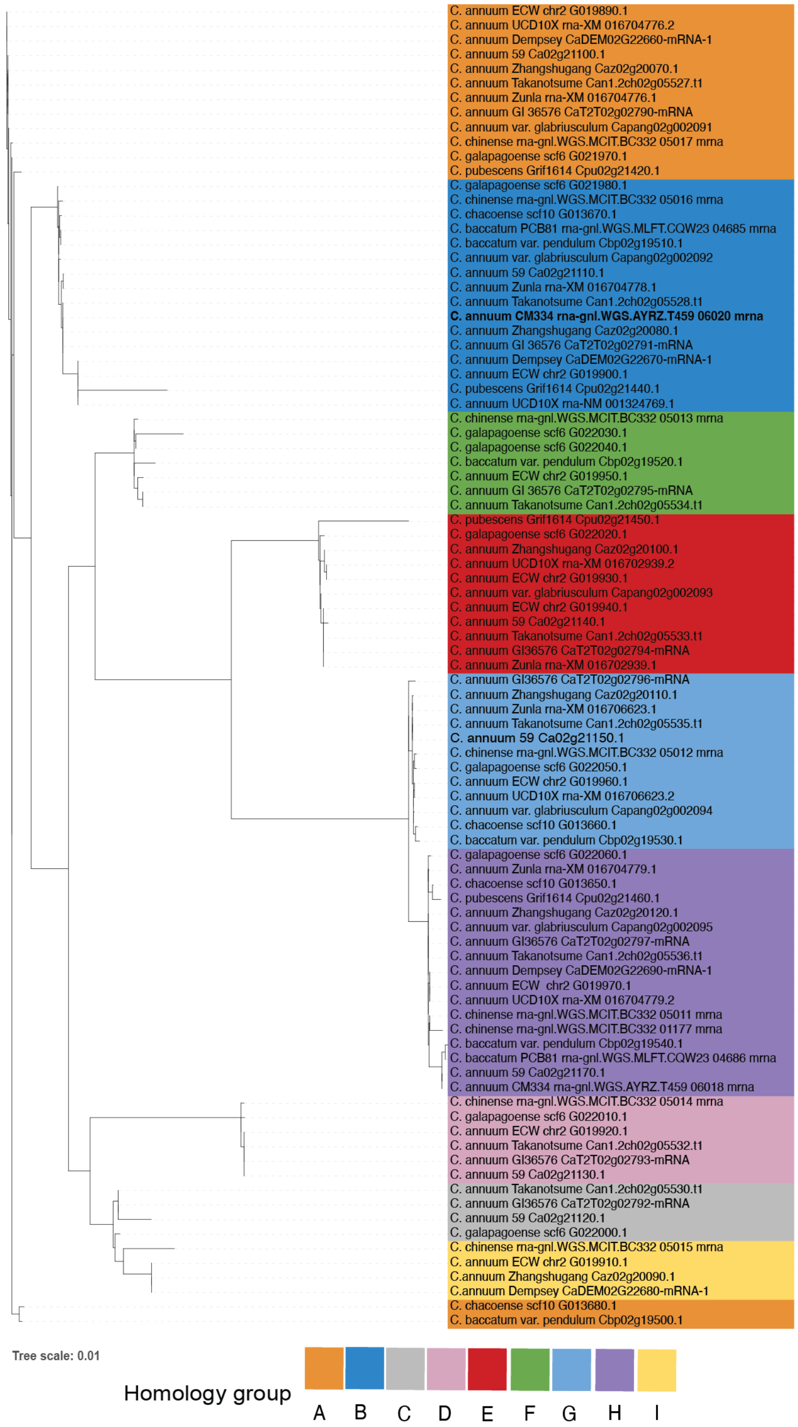


**Figure S14**: A Maximum likelihood tree for the members of the nine homology groups with a blast hit with the *Pun1* protein. The clustering of the sequences majorly confirms the homology grouping in the nine respective groups (shown in nine different colors) with group highlighted in dark blue containing the members of capsaicin synthase 2(*C. annuum* CM334 homolog in bold).
